# Scalable intracellular delivery via microfluidic vortex shedding enhances the function of chimeric antigen receptor T-cells

**DOI:** 10.1101/2024.06.25.600671

**Authors:** Brandon J. Sytsma, Vincent Allain, Struan Bourke, Fairuz Faizee, Mohsen Fathi, Rebecca Berdeaux, Leonardo M.R. Ferreira, W. Jared Brewer, Lian Li, Fong L. Pan, Allison G. Rothrock, William A. Nyberg, Zhongmei Li, Leah H. Wilson, Justin Eyquem, Ryan S. Pawell

## Abstract

Adoptive chimeric antigen receptor T-cell (CAR-T) therapy is transformative and approved for hematologic malignancies. It is also being developed for the treatment of solid tumors, autoimmune disorders, heart disease, and aging. Despite unprecedented clinical outcomes, CAR-T and other engineered cell therapies face a variety of manufacturing and safety challenges. Traditional methods, such as lentivirus transduction and electroporation, result in random integration or cause significant cellular damage, which can limit the safety and efficacy of engineered cell therapies. We present hydroporation as a gentle and effective alternative for intracellular delivery. Hydroporation resulted in 1.7- to 2-fold higher CAR-T yields compared to electroporation with superior cell viability and recovery. Hydroporated cells exhibited rapid proliferation, robust target cell lysis, and increased pro-inflammatory and regulatory cytokine secretion in addition to improved CAR-T yield by day 5 post-transfection. We demonstrate that scaled-up hydroporation can process 5 × 10^8^ cells in less than 10 s, showcasing the platform as a viable solution for high-yield CAR-T manufacturing with the potential for improved therapeutic outcomes.

## INTRODUCTION

Adoptive chimeric antigen receptor (CAR) T-cell therapy requires the *ex vivo* modification of a donor’s T-cells to express an engineered surface receptor, the CAR, that recognizes a unique tumor antigen *via* a single-chain variable fragment (scFv). Typically, this transmembrane receptor is linked to intracellular signaling domains like CD3ζ paired with either 4-1BB or CD28 co-stimulatory domains^1,2^. After the stable gene modification has been conferred, CAR-Ts are expanded and then re-infused into the patient, where they specifically target tumor cells for lysis. Recent clinical studies have demonstrated the efficacy of anti-CD19 CAR-T therapy, with complete remission being reported in patients with acute or chronic lymphoblastic leukemias and B-cell lymphomas^1,3,4^. This led the U.S. Food and Drug Administration (FDA) to approve multiple CD19-directed CAR-T therapies for B-cell malignancies^5,6^. Despite the curative potential of these therapies, there are still major safety concerns regarding the toxicity associated with the powerful immune effector response induced by robust CAR-T activation, such as cytokine release syndrome (CRS) and immune effector cell-associated neurotoxicity syndrome (ICANS)^7,8^.

These clinical results show the enormous potential of CAR-T therapy as a treatment for hematologic malignancies. However, there are still several hurdles to overcome before its adoption as a first-line treatment for many cancer types. Current manufacturing methods rely on lenti- or retroviral transduction for stable integration of the CAR transgene into the T-cell genome^9^. While this is a highly efficient approach, it lacks the precision editing that avoids insertional mutagenesis caused by random integration into the genome, potentially leading to oncogenic and mutagenic CAR-T products^10,11^. This raises serious safety concerns for clinical applications^12–14^. Furthermore, the strict regulations mandated for current Good Manufacturing Practice (cGMP) for viral production laboratories make the process slow and expensive, which limits its feasibility for clinical- and commercial-scale manufacturing.

Precision genome editing overcomes these safety concerns by specifically integrating the transgene into a defined locus within the host genome. For example, clustered regularly interspaced short palindromic repeats (CRISPR)-CRISPR-associated protein 9 (Cas9) CRISPR/Cas9 is a targeted nuclease that can induce a double stranded break (DSB) at a defined location^15,16^. By providing donor template DNA with homology arms complementary to the sequences that flank the cut site, the homology directed repair (HDR) pathway can then integrate the donor DNA sequence into that defined location, acting in a dual capacity to both knockout the gene of interest, and knock-in your payload of choice.^17,18^. In light of these advances, a series of seminal papers have been published which demonstrate the utility of CRISPR/Cas9 as a tool for adoptive T-cell engineering^15,16,19–22^. Generally, these approaches utilize electroporation, or nucleofection, as a mechanism for intracellular delivery of gene editing payloads, where the cell membrane is permeabilized by exposing cells to one or more electric pulses of varying amplitude and duration. This creates pores in the plasma membrane by which the payload can enter the cell^23,24^. As demonstrated by Pal *et al*., 2024., use of a ribonucleoprotein (RNP) and non-integrating adeno-associated viruses (AAV) to generate CAR-Ts holds significant promise in clinical trials, particularly in treatment of clear cell renal cell carcinoma (ccRCC)^25^. The authors showed that CD70-targeting CAR-T therapy resulted in complete regression of ccRCC xenograft tumors in a mouse model and disease control in 81.3% of patients in the clinic^25^. CRISPR engineered CAR-T cells have also been used against CD19 for hematological malignancies. Stadtmauer *et al*., 2020, demonstrated in their Phase 1 pilot the safety and feasibility of using multiplex CRISPR-Cas9 T-cells^26^. Zhang *et al.,* 2022, used non-viral delivery to develop anti-CD19 CAR-T cells, integrated into the *PD1* gene, observing a high rate (87.5%) of complete remission and durable response in 8 patients with relapsed/refractory aggressive B cell non-Hodgkin lymphoma^27^.

Using electric pulses to permeabilize the plasma membrane remains one of the most common intracellular delivery methods to date. Both the Neon Transfection System (electroporation) and the Lonza 4D-Nucleofector (nucleofection) have been adopted as industry standards for intracellular delivery, particularly with hard-to-transfect cell types such as primary human T cells, since there exist unique buffer solutions and pulsing protocols that are optimized for specific cell types^28,29^. While electroporation and nucleofection are efficient intracellular delivery mechanisms, there are severe drawbacks to this approach due to the significant damage caused by the electrical pulse^30^. Firstly, intracellular membrane-bound organelles are targeted by these pulses in addition to the plasma membrane. This results in the leakage of destructive enzymes from lysosomes, pro-apoptotic factors from mitochondria, and various cytoplasmic components out of the cell^31,32^. Furthermore, electroporation has been shown to potentially cause irreversible genomic DNA damage as well as produce reactive oxygen species (ROS), which cause further damage to DNA, proteins, and lipids within the cell^33–36^. Together, these effects put immense strain on electroporated cells, which can cause a high degree of cell death.

Herein, we describe hydroporation as an alternative intracellular delivery mechanism amenable to CAR-T generation for research and clinical applications. Hydroporation employs microfluidic vortex shedding (*µVS*), a hydrodynamic phenomenon whereby oscillating fluid forces gently permeabilize the cell membrane, allowing delivery of gene editing payloads such as Cas9 ribonucleoprotein (RNP) complexes.

Hydroporation relies on posts spaced approximately twice the typical cell diameter, resulting in cell-size independent delivery and flexible throughput (ie. 10^4^∼10^8^ activated T-cells mm^-1^ flow cell width) in a manner that is gentler than electroporation. Additionally, hydroporation utilizes 10-50x less reagents (*i.e.*, mRNA, RNP) per cell than other nascent delivery platforms^37–40^. These fundamental characteristics of hydroporation lend themselves to cell therapy manufacturing where starting material between donors and patients can vary significantly, perturbation adversely affects cell function, and a significant number of cells need to be processed^41,42^. We have previously demonstrated the utility of hydroporation as a mechanism for intracellular delivery of mRNA and Cas9 RNP targeting the *TRAC* locus in primary T cells^43,44^. Here, we target *TRAC* disruption, through RNP delivery via hydroporation, with an anti-CD19 CAR AAV homology directed repair template (HDRT) to achieve precision editing of CAR-Ts (Fig. 1A)^16,45^.

**Fig. 1.**
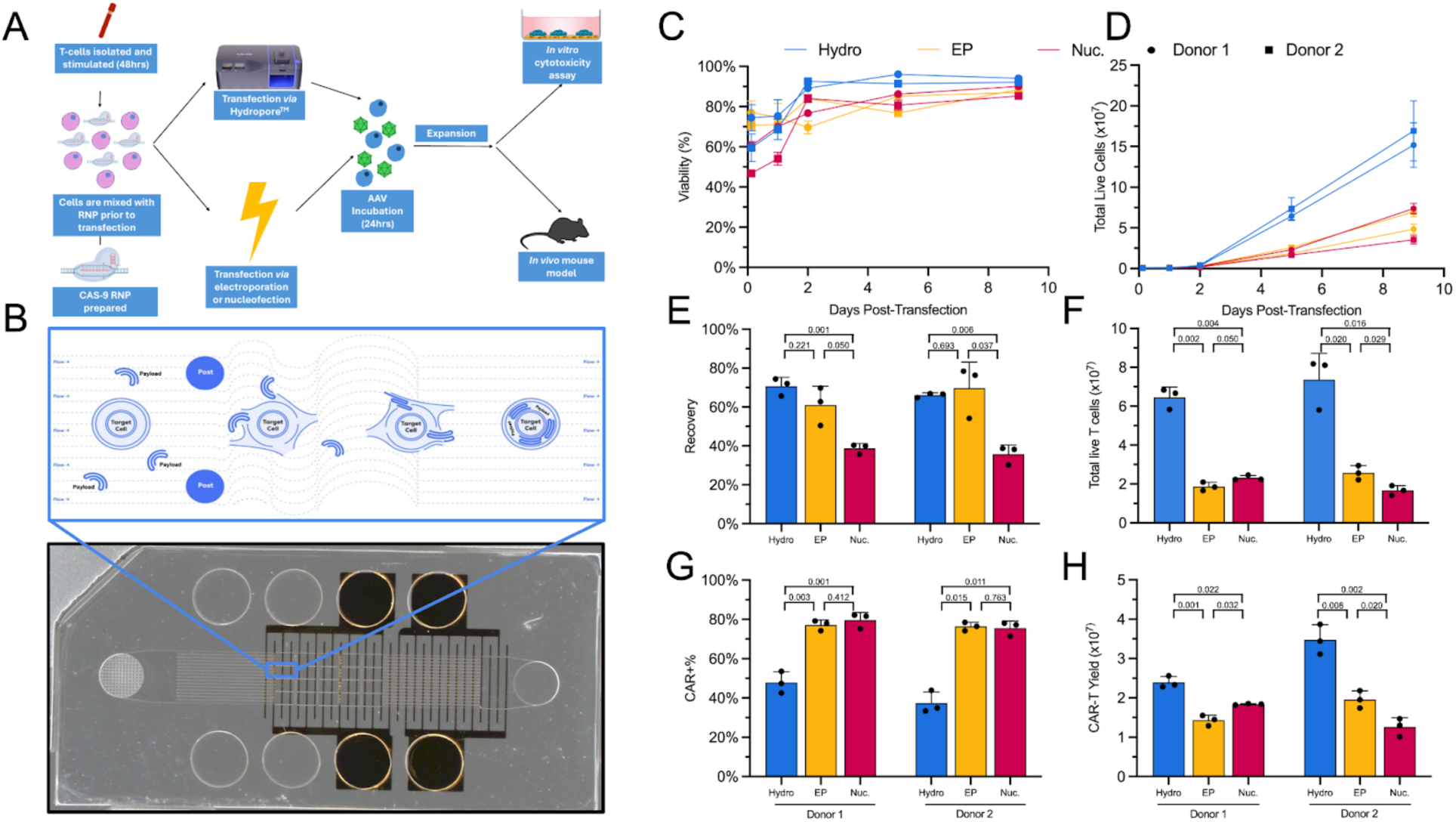
CAR-Ts transfected via hydroporation showed similar viability, proliferation, and EGFR+ cells, but improved CAR-T yield, when compared to electroporated or nucleofected cells. **A)** Overview of workflow for generation of CAR-Ts through RNP and AAV transfection, and how hydroporation is incorporated into the cell therapy workflow with improved cell numbers and viabilities for downstream analysis. **B)** Picture of the microfluidic hydroporation chip, with an illustration of the microfluidic vortex shedding that cells undergo within the chip (blue window). **C)** Head-to-head viability comparison of 2 sets of T-cell donors transfected by hydroporation (Hydro; Blue), electroporation (EP; Yellow) or nucleofection (Nuc; Magenta). **D)** Proliferation of T-cells from day 0 (transfection) to day 9. **E)** Recovery of T-cells 2 hours post transfection. **F)** Total live T-cells on day 5. **G),** Percentage of CAR+ T-cells, based on EGFR signal. **H)** Total CAR-T yield on day 5. All data points involve n = 3 technical replicates and, where relevant, p-values from two-tailed heteroscedastic unpaired t-tests.

In comparing the performance of hydroporation to nucleofection and electroporation in generating CAR-Ts, we evaluated the transfection efficiency, viability, proliferation and total CAR-T yield. Hydroporation resulted in 1.7- to 2-fold greater yield of CAR-Ts on average by day 5 post-transfection compared to electroporation and nucleofection, respectively. To further demonstrate the versatility of hydroporation, we designed a high-throughput microfluidic chip capable of processing up to 5 × 10^8^ activated T-cells in less than 10 seconds, showing that hydroporation is scalable and capable of processing a wide range of cell numbers with similar yields in recovery, viability, and editing efficiency.

Next, we sought to characterize the T-cell subset and cytokine secretion profile of CAR-Ts. Hydroporated CAR-Ts (Hy-CAR-Ts) and nucleofected CAR-Ts (Nuc-CAR-Ts) displayed nearly identical CD4+/CD8+ T-cell ratios, as well as naive/memory phenotype ratios. Both Hy-CAR-Ts and Nuc-CAR-Ts demonstrated specific activation and pro-inflammatory cytokine secretion upon target cell engagement.

Finally, in order to evaluate the potency and clinical utility of Hy-CAR-Ts, we assessed their functionality using *in vitro* and *in vivo* models that mimic recent successful clinical trials utilizing AAV-mediated CAR-Ts, and compared them to Nuc-CAR-Ts^25^. We observed similar killing potential for Hy-CAR-Ts, though Hy-CAR-Ts showed lower levels of activation-induced cell death (AICD) and improved motility in *in vitro* single cell serial killing assays. In addition, we observed equivalent *in vivo* activity between Hy-CAR-Ts and Nuc-CAR-Ts in an aggressive mouse xenograft model of human B-cell acute leukemia.

Thus, hydroporation offers a means of generating high yields of highly functional precisely genome-edited CAR+ T cells, reducing manufacturing costs and time needed to generate engineered cell therapies like CAR-T.

## RESULTS

We have previously demonstrated the utility of hydroporation (i.e. *µVS*) as a mechanism for intracellular delivery of both mRNA and Cas9 RNP to primary human T-cells^43,44^. Importantly, previous studies indicated minimal dysregulation of the native T-cell state along with rapid cell recovery and proliferation following hydroporation. In order to evaluate the potential of hydroporation as a tool for adoptive immunotherapy manufacturing, we compared hydroporation with nucleofection and electroporation in the generation of CAR-Ts. Using a previously validated CRISPR knock-in system, we utilized Cas9 RNP targeting the first exon of *TRAC* and an AAV vector encoding a self-cleaving P2A peptide, upstream from the HDR template for TRAC, consisting of a CD19-specific 1928z CAR and truncated epidermal growth factor receptor (EGFRt) reporter^16^. Hydroporation, nucleofection or electroporation of activated primary human CD3+ T cells took place 2 days after being isolated from frozen peripheral blood mononuclear cells (PBMCs). Cells were cultured for 9 days following RNP and AAV delivery and were evaluated for viability, proliferation, knock-out (KO) efficiency, knock-in (KI) efficiency, and CAR-T yield (Fig. 1).

The fluid dynamic conditions created in the post array region of the microfluidic chip gently and efficiently permeabilize the plasma membrane to promote external material uptake (Fig. 1b and Supplemental Fig. 1). Compared to electroporation and nucleofection, this method of membrane poration is less detrimental to cell health, which is reflected in the improved recovery and viability in hydroporated cells (Fig. 1c and 1e). Indeed, hydroporated cells reach >90% viability within 2 days following transfection, while nucleofected and electroporated cells only reach 90% viability at 5 days post-transfection. In terms of cell recovery, hydroporation and electroporation performed similarly, with 60-70% live cell recovery 2 hours after RNP delivery. Nucleofection showed the lowest recovery, with <40% live cell recovery.

Both nucleofection and electroporation resulted in highly efficient KI rates (∼80%), while hydroporation-mediated transgene insertion was lower, at ∼40%, though the KI:KO ratio was identical for all methods with 80% of KOs converting to KIs in the presence of the HDRT. (Fig. 1g and Supplemental Fig. 2). However, in the 5 days following payload delivery, hydroporated cells divided more rapidly, resulting in 3.2x and 3.6x more live cells than nucleofection or electroporation on average, respectively (Fig. 1d and f). When taken with the corresponding recoveries and KI efficiencies, hydroporation yielded 1.7x and 2.0x more CAR-Ts, on average, than electroporation and nucleofection (Fig. 1h).

Unlike static electric pulsing techniques, such as nucleofection or electroporation, which use cuvettes or pipette tips equipped with electrodes, hydroporation employs a stable flow delivery model which lends itself to be easily scaled up or down depending on the desired transfection volume (Fig. 2a). Here, we demonstrated the versatility of hydroporation by designing arrayed versions of our proprietary flow cell in order to accommodate greater numbers of cells. For small scale transfections (<10^8^ cells) we used chips containing either 1 flow cell or 4 sub-flow cells which process cells simultaneously, known as Research Use Only (RUO) chips. For large scale transfections aimed at manufacturing a clinically relevant dose of CAR-Ts, we employed microfluidic chips which contained 40 arrayed flow cells, known as Cell Therapy (CT) chips.

**Fig. 2.**
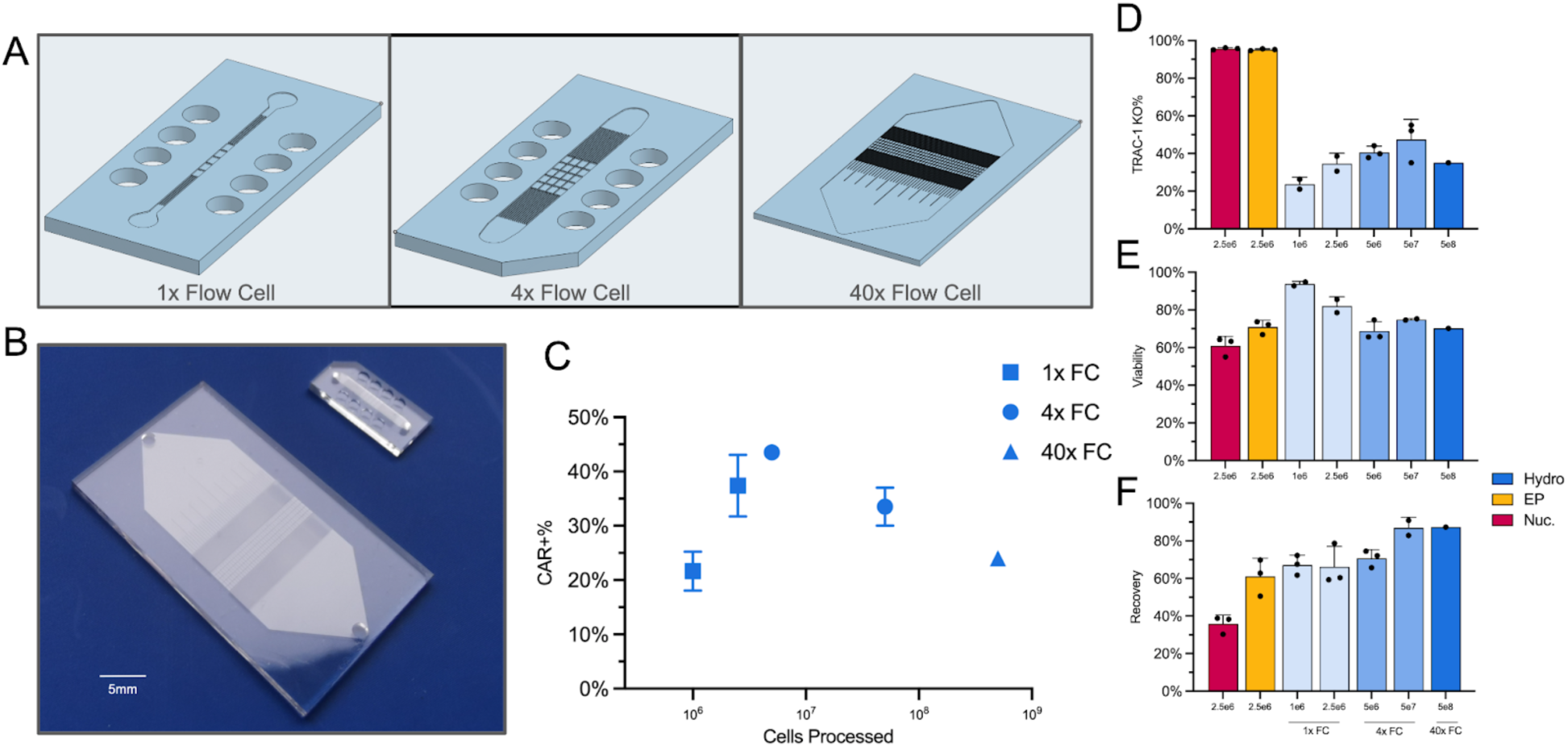
Scaled up version of the hydroporation chip, shows similar TRAC-1 KO, viability and recovery when compared to electroporation and nucleofection even when processing 1 billion activated T-cells. **A)** CAD images of the small volume RUO chip (1x flow cell), standard volume RUO chip (4x Flow cell) and large volume CT chip (40x Flow cell), with an enlarged image showing the post design per flow cell. **B)** Physical representation of CT chip (40x flow cells, left) to RUO chip (1-4x flow cells, right), bar 5 mm. **C)** Percentage of CAR+ T-cells on day 5 after transfection of 10^6^ - 5 × 10^8^ total cells. **D)** TRAC KO% efficiency, against cell numbers, based on whether cells were hydroporated (Hydro; blue), electroporated (EP; yellow) or nucleofected (Nuc, Magenta). **E)** TRAC KO Cell viability 24 hours post-transfection. **F)** Cell recovery 2 hours post transfection. All data points involve n = 3 biological replicates except 40x FC data.

When different numbers of cells are processed, there is a peak of around 40% CAR+ T cells generated at 10^7^ T cells with the 4x flow cell RUO chip (Fig. 2b), though there is no significant difference when processing higher cell numbers (10^9^ cells showed around 25% CAR+ T-cell efficiency). There is a trend where the greater the number of processed cells, the higher the % of *TRAC* KO (Fig. 2c), particularly going up to 5 × 10^6^ cells in the 4x flow cell chip (see Figure 2). Cells at this density also display approximately 70% viability (Fig. 2d), similar to that observed with electroporated and nucleofected cells. This trend is also observed in cell recovery, with hydroporation allowing for the recovery of a higher number of transfected cells compared to both nucleofection and electroporation (Fig. 2e). For hydroporation, higher cell numbers equated to greater recovery and transfection efficiency. Total sample processing times, for both the RUO and CT chips, range from milliseconds to seconds. For example, the RUO chip processes a typical sample of 5 × 10^6^ activated T-cells (*i.e.,* 100 µL at 5 × 10^7^ cells mL^-1^) in less than 1 second, while the CT chip can handle 5 × 10^8^ activated T-cells (*i.e.*, 5 mL at 10^8^ cells mL^-1^) in under 10 seconds.

To further understand the impact of the transfection method on T-cell phenotype, the proportions of naive and memory cell subsets in CD4^+^ and CD8^+^ (CAR) T cells were determined by flow cytometry for both Hy-CAR-Ts and Nuc-CAR-Ts (Supplemental Fig. 3). We did not observe any significant differences in the frequencies of these subpopulations in CD4^+^ or CD8^+^ T cells from either donor between Hy-CAR-Ts and Nuc-CAR-Ts (Fig. 3a).

**Fig. 3.**
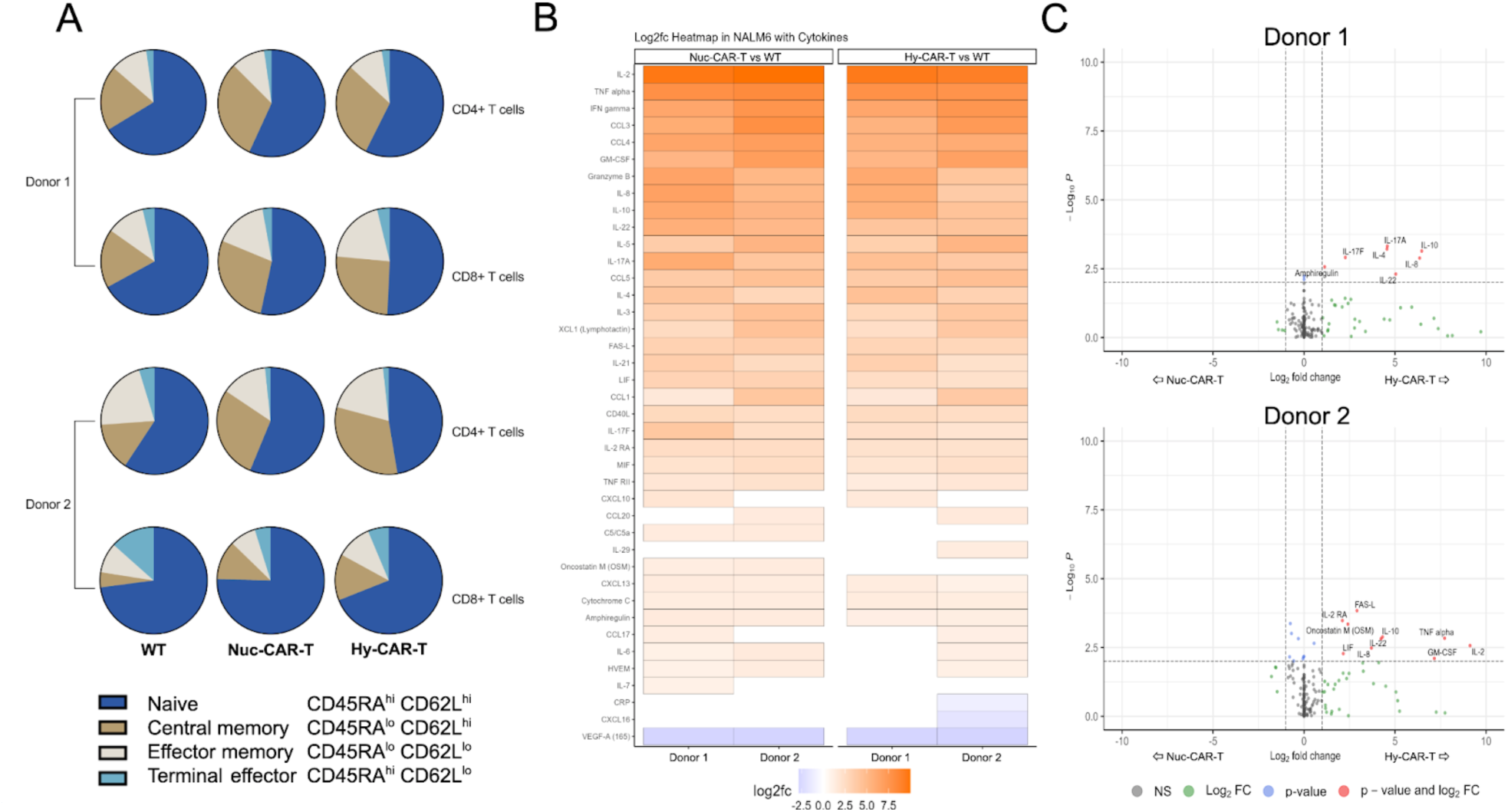
T-cell phenotype and cytokine profile show no significant differences between hydroporation and nucleofection. **A)** Relative proportions of naive/memory CD4^+^ and CD8^+^ T-cell subsets for CAR+ cells engineered via hydropore (Hy-CAR-T) or nucleofection (Nuc-CAR-T). Percentages of naive/memory cells for both CD4^+^ and CD8^+^ T-cells were calculated based on expression of CD45RA and CD62L. **B)** Heatmap of 40 proteins secreted by CAR-T cells that were significantly differentially expressed compared to WT cells in the presence of NALM6 target cells. **C)** Comparison of cytokine secretion patterns of CAR-T cells from donor 1 and donor 2 generated using either nucleofection or hydroporation upon activation with NALM6 target cells at a 1:1 ratio, as assessed by 187-plex nELISA. NS: not significant; FC: fold-change.

To assess the function of Hy-CAR-Ts, we first characterized the secreted cytokine profile of the cells upon engagement with target NALM6, a CD19-expressing B-cell leukemia cell line, using nELISA. For this study, we co-cultured Hy-CAR-Ts, Nuc-CAR-Ts or wild type (WT) T cells with NALM6 B-cell leukemia target cells overnight at a 1:1 ratio. The next day, supernatants were collected for protein quantification. For both donors tested, Hy-CAR-Ts and Nuc-CAR-Ts secreted high levels of pro-inflammatory cytokines, such as IL-2, IFNɣ, TNFa and IL-17A along with chemokines, such as IL-8, CCL1, CCL5, and the cytotoxicity effector molecule granzyme B, compared to WT samples (Fig. 3b). These cytokine and chemokine expression patterns indicate robust activation via the anti-CD19 CAR.

We also compared the cytokine secretome of Hy-CAR-Ts directly with Nuc-CAR-Ts. While the number and magnitude of significantly different cytokines was relatively small, Hy-CAR-Ts displayed higher levels of cytokine secretion in both donors analyzed (Fig. 3c). Interestingly, between the donors involved in this study, there was a common upregulated anti-inflammatory cytokine in activated Hy-CAR-Ts compared with Nuc-CAR-Ts - IL-10. Moreover, the pro-inflammatory chemokine IL-8 and the IL-10 family cytokine IL-22 were also upregulated in Hy-CAR-Ts compared with Nuc-CAR-Ts in both donors.

T-cell populations are highly variable, and cell phenotypes change after interaction with target cells. Therefore, we undertook a single-cell approach, Time-lapse Imaging Microscopy In Nanowell Grids (TIMING™)^46,47^, to compare the dynamics and functions of Hy-CAR-Ts with Nuc-CAR-Ts from each of five different donors (Fig. 4). Cells are constrained within nanowells, allowing identification and tracking of individual cells over time before, during, and after interactions with other cells (Fig. 4a and Supplemental Video 1).

**Fig. 4.**
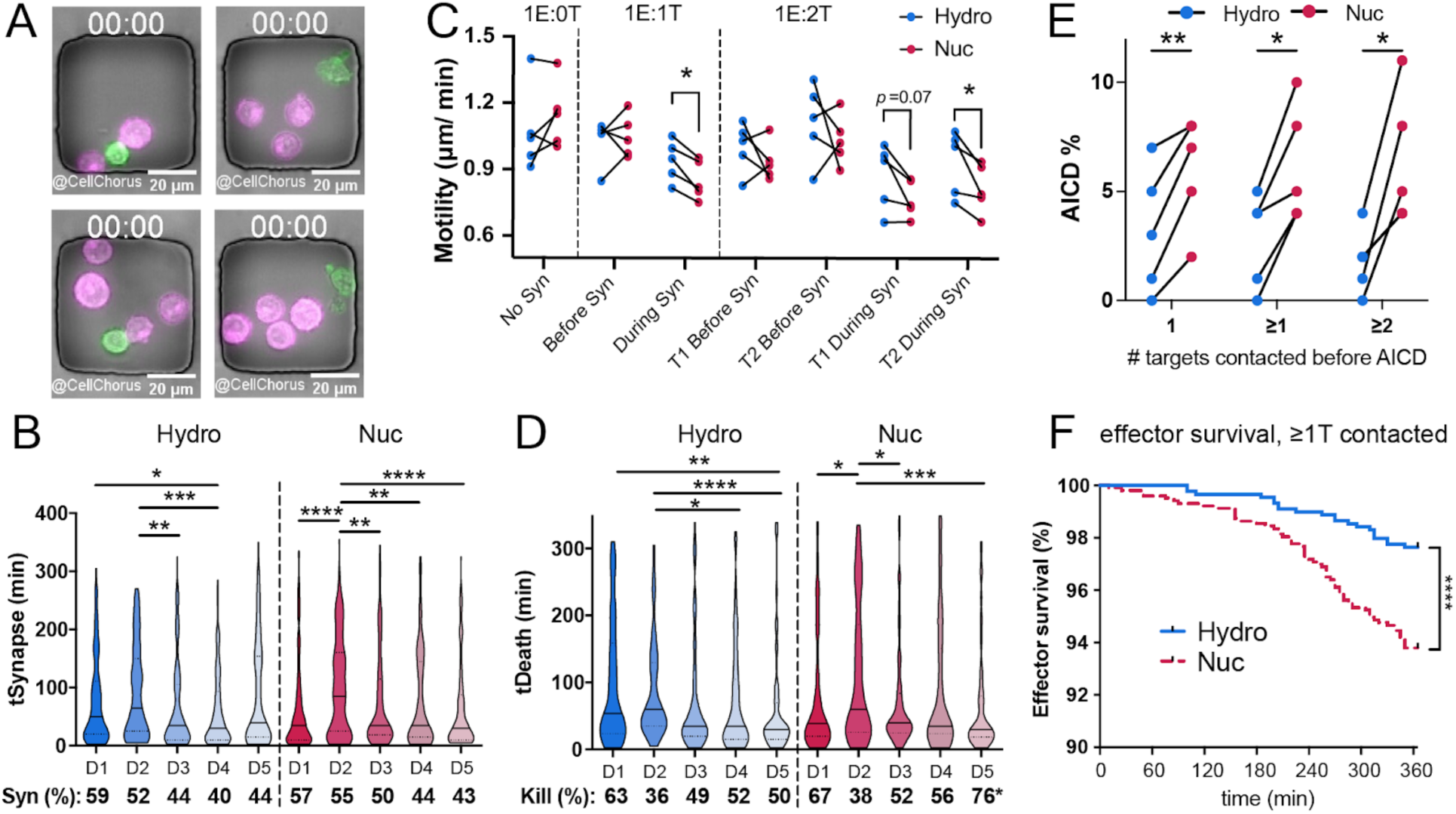
CAR-T cells manufactured with hydroporation have superior motility and resistance to AICD. **A)** Images of cells in TIMING nanowells at different E:T ratios at the beginning of TIMING assays (00:00 hh:mm). CAR-T effectors (green), NALM6 target cells (magenta), bars 20 µm. **B)** Violin plots show population dynamics of synapse duration (tSynapse) between Hy-CAR-T (blue) or Nuc-Car-T (magenta) and NALM6. Bottom line indicates percent of CAR-T observed to form a synapse (Syn) in each assay. **C)** CAR-T motility (µm/ min displacement) before or during synapse with target cells. Each point is the mean for each donor (D1-D5), shown as paired samples of Hy-CAR-T (blue) vs Nuc-CAR-T (magenta). *p<0.05 by pairwise Mann-Whitney tests. **D)** Time required for CAR-T cells to kill target cells (tDeath of target, min) after synapse formation. Bottom line shows percent of synapses resulting in a kill. tDeath distributions: *p<.05, **p<0.01, ***p<0.001, ****p<.0001 by pairwise Mann-Whitney. Kill % difference: *p < .05 by Fisher’s exact test on raw numbers (only D5 showed differential killing of Hy-CAR-T vs Nuc-CAR-T). **E)** Mean AICD of effectors after engagement with one or more target cells (x-axis). Each connected pair of points indicates an individual donor. **p<.05, **p<0.01 by 2-tailed t-tests. F) Survival curves of effectors after target contact (AICD) over 6-hour TIMING assay. Curves represent pooled data from three donors (D3-D5). ****p<.0001 by Kaplan-Meier analysis.

TIMING revealed differences between Hy-CAR-T and Nuc-CAR-T that cannot be detected in bulk assays^47^. We observed intrinsic donor variability in CAR-T target seeking and contact dynamics, but no significant differences in these parameters between CAR-T prepared by hydroporation and nucleofection. For example, the time required for CAR-T cells to form a stable synapse with target cells (“tSeek”, Supplemental Fig. 4a) and individual synapse duration (“tSynapse”) with target cells varied among donors, but there were no differences between the matched Hy-CAR-T and Nuc-CAR-T prepared from the same donor (Fig. 4b). During target engagement (synapse), Hy-CAR-Ts derived from all five donors maintained significantly more motility than Nuc-CAR-T cells derived from the same donor (Fig. 4c), suggesting a potential benefit for Hy-CAR-Ts based on prior data linking single-cell motility with resistance to exhaustion^48,49^.

Notably, the difference in CAR-T cell motility was not observed in the absence of target cells or prior to synapse formation. We observed no significant differences between Hy-CAR-T and Nuc-CAR-T cells in the percent of CAR-T cells able to form a synapse, kill, or serial kill (Supplemental Fig. 4a, b) or the time required to kill targets, though there was substantial donor variability (Fig. 4d).

A major advantage of TIMING is that, by constraining cells in nanowells, the fate of effectors after target killing is quantifiable^50^. Consistent with lower motility during synapse, we found that Nuc-CAR-T were more sensitive to AICD than Hy-CAR-T cells from the same donor (Fig. 4e). AICD occurred more frequently in Nuc-CAR-T irrespective of the number of targets engaged, but AICD in Nuc-CAR-T tended to increase with an increased number of targets engaged. The difference in AICD was also significant when survival times were compared by Kaplan-Meier analysis of three pooled donors (Fig. 4f). Interestingly, we observed no significant differences in effector survival in nanowells containing only one effector and no target cells (Supplemental Fig. 4g), indicating that the increased cell death observed in Nuc-CAR-T is not due to the health of the cells before activation. Together, single-cell results suggest that CAR-T prepared by hydroporation might be more likely to resist exhaustion *in vivo*.

Encouraged by the above *in vitro* results, we next sought to compare the cytotoxicity of Hy-CAR-Ts with Nuc-CAR-Ts using an *in vitro* co-culture assay with CD19-expressing NALM6 cells (Fig. 5a) and a similar *in vivo* potency study (Fig. 5c,d). To enrich the CAR+ cell population to reduce noise in these studies, both Hy-CAR-Ts and Nuc-CAR-Ts were subject to TCR (T-cell receptor) depletion. Hy-CAR-Ts demonstrated similar target cell lysis at all E:T ratios when compared with Nuc-CAR-Ts in this bulk killing assay (Fig. 5b). In addition to the *in vitro* co-culture assay, an *in vivo* ‘stress test’, in which the CAR-T dose is lowered to reveal the functional limits of different CAR-T populations, was performed as previously reported^45^. A total of 10^5^ or 4 × 10^5^ CAR^+^ T-cells, generated using either hydroporation or nucleofection, were injected intravenously into immunodeficient NOD.Cg-*Prkdc^scid^ Il2rg^tm1Wjl^*/SzJ (NSG) mice engrafted with 5 × 10^5^ NALM6 cells 4 days prior. An assessment on day 100 indicated that the low dose of Hy-CAR-T and Nu-CAR-T may improve survival relative to the high dose for both donors, however, a longer study may be required given that >40% of mice survived to 100 days in all CAR-T conditions (Fig. 5d). Tumor burden was evaluated over time by bioluminescence imaging (BLI), indicating a similar tumor burden for both transfection methods at either CAR-T dose (Fig. 5c and Supplemental Fig. 5). Taken together, these results demonstrate that Hy-CAR-Ts have similar therapeutic potency to their electroporation counterpart, which has been previously observed in a similar study comparing electroporation^45^.

**Fig. 5.**
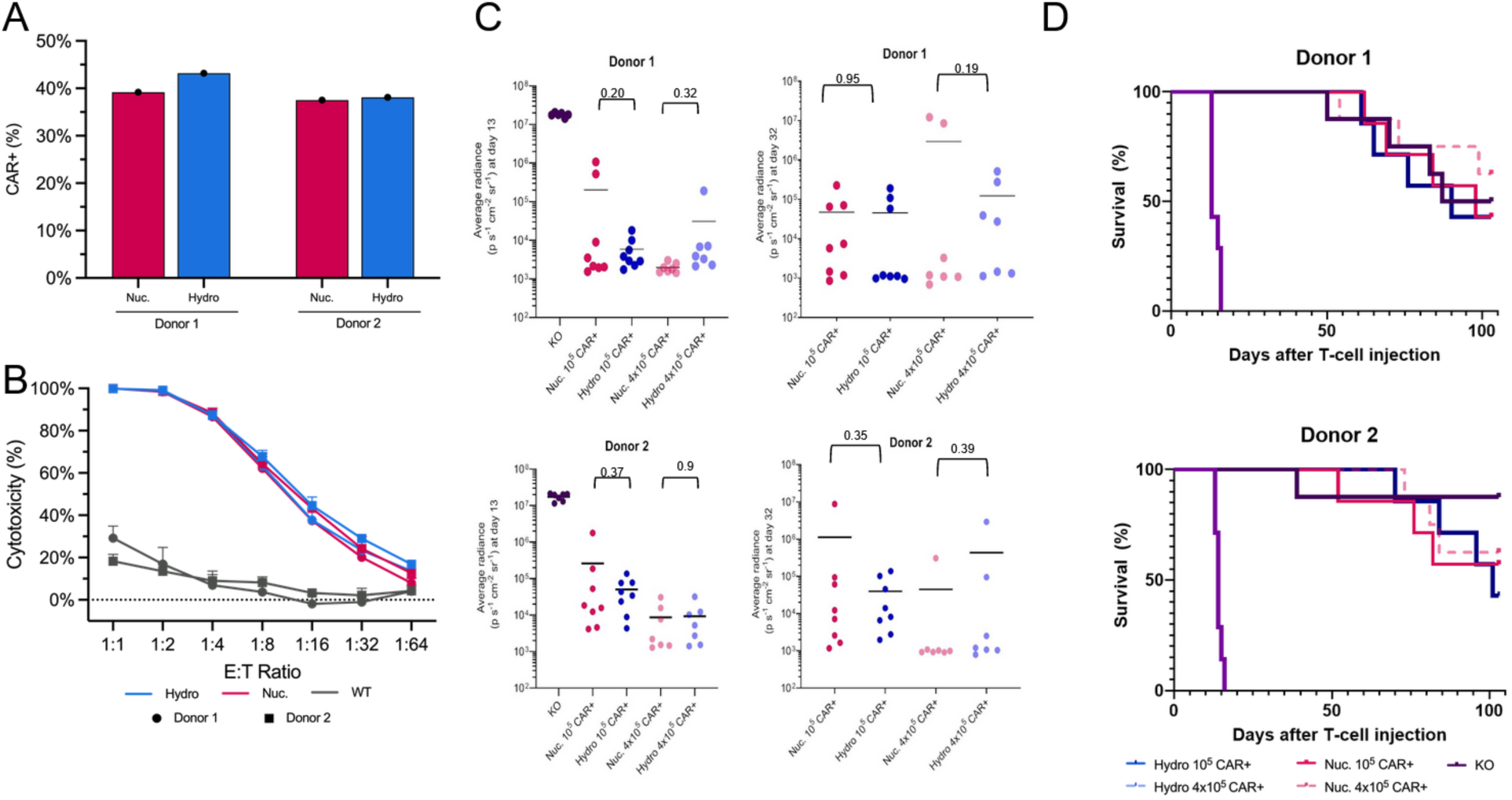
CAR-Ts modified through hydroporation showed similar yields of CAR+ cells as nucleofected T-cells, as well as similar potency in vivo as shown in survival rates of mice at Day 100. **A)** CAR-T+% after TCR depletion for donors 1 and 2 transfected by hydroporation or nucleofection. **C)** Bulk cytotoxicity assay, based on treated T-cells at different ratios of effectors to target cells (CAR-T to NALM6). **C)** BLI values, for donors 1 and 2, on the last measurement day on which all mice were alive (Day 13) and which all CAR-T injected mice were alive (Day 32). P values are from two-tailed Welch’s unpaired t-tests. **D)** Kaplan–Meier survival analysis of mice treated with/without hydroporated or nucleofected CAR+ T-cells, over 100 days. N = 8 mice per group.

## DISCUSSION

The delivery of gene editing payloads to primary human T-cells continues to be an obstacle in the development of robust manufacturing methods for adoptive immunotherapy. Lenti- and retro-viral gene modification have severe limitations, including cost, safety, and scalability. Furthermore, gene delivery platforms that utilize electroporation or nucleofection have demonstrated a significant negative impact on cell health and function, which limits their potential for clinical applications. Recent studies that use Cas9-RNP show increasing utility in the specific integration of CARs or TCRs into the *TRAC* locus^52,53^. Other intracellular delivery methods exist^54^, including peptide-enabled RNP delivery for CRISPR (PERC)^45^, microneedle injection^37^, and cell squeezing^39^. However, these methods typically suffer from drawbacks like low throughput, cell size-dependent delivery, high reagent consumption, and user safety concerns^55^.

These challenges motivated us to investigate microfluidic vortex shedding as a mechanism for cell membrane permeabilization and subsequent intracellular delivery of gene editing payloads. The advantages of the microfluidic transfection platform are evident in its gentle nature, ease of use, scalability, and reduced cost. Prior studies of hydroporation using mRNA^43^ and Cas9 RNP^44^ showed the potential for this platform as a tool for T-cell engineering, though the advantages were not fully realized until we achieved KI of an anti-CD19 CAR into the *TRAC* locus. When compared to nucleofection, not only did hydroporation yield a significantly greater number of CAR-Ts, but those CAR-Ts exhibited equivalent or better function in the presence of target cells.

To better understand the dynamics of enhanced motility and reduced AICD or improved serial killing *in vitro* assessed with the TIMING assay, we analyzed the T-cell phenotype and cytokine production. There was no difference in the composition of the naive/memory T-cell subsets between Hy-CAR-Ts and Nuc-CAR-Ts. Of the 187 cytokines tested, 3 showed significant difference between activated Hy-CAR-Ts and Nuc-CAR-Ts during killing for both donors: IL-8, IL-10, and IL-22.

IL-8 is a pro-inflammatory chemokine that attracts and activates neutrophils^56^. IL-8 production by tumor infiltrating activated CAR-T can thus increase neutrophil tumor infiltration. IL-8 is also secreted by several types of tumors, prompting CAR-T researchers to explore the influence of IL-8 and antitumor efficacy in solid tumors^57^. IL-22 is a pleiotropic cytokine, with reported pro-inflammatory and anti-inflammatory effects depending on the context^58^. In colorectal cancer, IL-22-producing infiltrating CD4+ and CD8+ T-cells were correlated with a better clinical outcome and increased infiltration of neutrophils, which in turn enhanced anti-tumor T-cell responses^59^. Moreover, CAR-Ts engineered to secrete IL-22 were found to be more cytotoxic towards head and neck squamous cell carcinoma^60^.

While the vast majority of cytokines produced upon T-cell activation are pro-inflammatory, IL-10 can act as a regulatory interleukin and suppress the immune response as a built-in negative feedback loop. IL-10 works by inhibiting the production of other pro-inflammatory cytokines, suppressing the antigen-presenting capacity of dendritic cells and macrophages, reducing the expression of major histocompatibility complex (MHC) class II and co-stimulatory molecules on antigen-presenting cells and promoting the differentiation of regulatory T-cells (Tregs)^61^. This suggests that even though Hy-CAR-Ts have an extremely robust pro-inflammatory cytokine response to CAR activation, there is still production of regulatory cytokines to limit excessive inflammation.

From an efficacy standpoint, the persistence of CAR-Ts *in vivo* is closely linked to positive clinical outcomes. CAR-T proliferation *in vivo* is fundamental to long term persistence. Studies indicate that electroporation limits the proliferative capacity of cells due to DNA damage as well as the generation of ROS caused by the electric pulse(s). As demonstrated by our results, hydroporated cells maintain a high proliferative capacity *in vitro* compared to nucleofected cells, which could potentially translate into improved persistence *in vivo*. CAR-T’s persistence *in vivo* is far more nuanced than simple proliferative capacity. Recent literature shows that the tumor microenvironment can be very inhospitable to CAR-Ts, leading to AICD and exhaustion^50,62,63^. Our results, though confined to a 6 hour window, show significantly lower levels of AICD with improved motility in CAR-Ts generated by hydroporation compared to nucleofection. This suggests that Hy-CAR-Ts are better equipped to robustly engage target cells without succumbing to AICD, which would potentially lead to longer persistence and a greater anti-tumor effect.

## CONCLUSIONS & FUTURE WORK

We successfully used microfluidic vortex shedding or hydroporation to generate genome edited CD19-directed CAR-Ts with superior proliferative capacity, improved motility and reduced AICD when compared to nucleofected CAR-Ts. Hy-CAR-Ts were then shown to have (1) increased IL-10 production upon CAR engagement – a critical regulatory cytokine for CAR-Ts, especially for solid tumors – all while (2) maintaining potency *in vivo* in a B-cell leukemia mouse xenograft model. We also demonstrated previously reported hydropore designs that maintain similar performance when scaled up and out to process 5 × 10^8^ activated T-cells in less than 10 seconds – a key criterion for cell therapy manufacturing.

Cumulatively, these results indicate that hydroporation can be incorporated into engineered cell therapy research and manufacturing processes due to: (1) flexible throughput; (2) cell-size independent delivery; (3) gentle processing; (4) improved yields due to higher proliferation; (5) reduced reagent consumption all (6) without loss of CAR-T function both *in vitro* and *in vivo*.

There are, however, still issues related to use of AAV for successful CAR KI due to size limitations. With this in mind, the hydroporation platform can be further developed to address these potential problems. Our future efforts are focused on multiplex editing and incorporating various forms of DNA to our KI strategies to exceed the size limitations of AAV. We see hydroporation being applied to other therapeutic, or difficult to transfect cell types, and we have already demonstrated the potential of hydroporation to knock-in a transgene in regulatory T cells (CAR-Tregs) and natural killer (NK) cell.

## METHODS

### Materials

#### Chip fabrication and operation

Chips were fabricated using previously reported designs^43^ and mask-based 3D printing (3D printed microtec, Bethesda, MD). Briefly, chips and chip features (*i.e.*, inlet filter, posts, channel thickness, inlet and outlet channels) were manufactured by (1) generating a digital rendering of the microfluidic chip using Onshape CAD software, (2) preparing a photomask for patterning GR-1 resin (pro3dure, audioprint® GR-1) and (3) performing a series of 3D printing, metallisation, bonding and inspection steps. Chips were operated with a hydroporation instrument using compressed nitrogen at 140 psig, as per previously reported protocols^44^.

#### Isolation & culture of primary human T-cells

Primary human CD3+ T cells were isolated by negative selection from cryopreserved PBMCs (STEMCELL Tech, CAT#70025) using the EasySep Human T-cell Isolation Kit (STEMCELL Tech, CAT#17951) per the manufacturer’s recommendation. After isolation, T-cells were seeded at 10^6^ cells mL^-1^ in complete culture media. Complete culture media consisted of X-VIVO 20 media (Lonza, CAT#04-44Q) supplemented with 5% human serum AB (Access Biologicals, CAT#535-HI) and sterilized through a 20 nm vacuum filter. T-cells were stimulated for 48 hours with bead conjugated CD3/CD28 Dynabeads at a 1:3 bead-to-cell ratio for *in vitro* studies and a 1:1 bead-to-cell ratio ahead of the *in vivo* study (Thermo Fisher, CAT#11132D). 48 hours after activation, Dynabeads are removed and T-cells are processed by Hydroporation, nucleofection or electroporation. Throughout the *in vitro* experiments, T-cells were maintained at a cell density of 10^6^ cells mL^-1^ and supplemented with 50 IU mL^-1^ rh IL-2 (Peprotech, CAT#200-02) every 2 days. Ahead of the *in vivo* study, cells were cultured in X-VIVO 15 media containing 5% human serum, 5 ng ml^−1^ IL-7 (Miltenyi, CAT#130-095-367) and 5 ng ml^−1^ IL-15 (Miltenyi, CAT#130-095-760).

#### RNP assembly

RNPs were produced by combining target-specific sgRNAs (Synthego) and recombinant Cas9 (Truecut V2, Thermo Fisher Scientific, CAT#A36499). Briefly, lyophilized sgRNAs were reconstituted in Nuclease-free molecular grade water (Invitrogen, CAT#AM9938) to a concentration of 100 µM and then aliquoted for storage at -20°C. On the day of transfection, sgRNAs were thawed and diluted to equal volume of Cas9 in nuclease free water, then mixed with Cas9 at a 1:1.5 Cas9 to sgRNA molar ratio. The RNPs were complexed by incubating the sgRNA:Cas9 mixture for 15 minutes at 37C, then moved to room temperature until used in the transfection (<1hr). 200 μg mL^-1^ RNP was then aliquoted into a clean eppendorf before adding cells for transfection *via* hydroporation, nucleofection or electroporation.

#### Transfection with RNP

48 hours after dynabead activation, T-cells were pelleted, washed with PBS, and thoroughly resuspended in OptiMEM (Gibco, CAT#31985062), P3 buffer with supplement (Lonza Bioscience, CAT#V4XP-3024), or R buffer (Thermo Fisher, CAT#MPK10025) at a cell density of 2 × 10^7^ cells mL^-1^. Before transfection, cells were added to Eppendorf tubes containing RNPs for a total volume of 100 µL, and mixed thoroughly via pipetting.

##### Nucleofection

For samples treated by nucleofection, following manufacturer guidelines, the 100 µL media consisting of cells, RNP in P3 buffer was added to 4D-Nucleofector cuvettes and treated using pulse code EH115. 400 µL of complete culture media was added to each cuvette and then transferred to a tissue culture incubator for 15 minutes for cell recovery. After the recovery period, Nucleofected cells were seeded at 10^6^ cells mL^-1^ in pre-warmed complete culture media and returned to the tissue culture incubator. For the *in vivo* study, cells were nucleofected in 20 µL reactions in the 96-well Lonza shuttle systems. Cell and payload concentrations were kept constant.

##### Electroporation

For samples treated by electroporation, following manufacturer guidelines, electrolyte E2 buffer was added to the Neon^®^ tube, and allowed to come to room temperature . Using the 100 µL Neon^®^ electroporation tip, the 100 µL media consisting of cells, RNP in R buffer was transferred to the Neon^®^ tube containing E2 buffer and treated at 1600 V, 10 ms and 3 pulses. Treated cells were transferred to pre-warmed complete culture media and seeded at 10^6^ cells mL^-1^ before being returned to the tissue culture incubator.

##### Hydroporation

For *in vitro* experiments, hydroporation was performed at 160 psi and 28.3V (2.25 kV cm^-1^). For *in vivo* experiments, hydroporation was performed at 140 psi and 0V. Hydroporated samples were collected in 15 mL conical tubes at 10^6^ cells mL^-1^ in pre-warmed complete culture media, then transferred to cultureware and kept in the tissue culture incubator.

In some cases, Nedisertib (M3814, MedChemExpress, CAT#HY-101570) was used at the manufacturer’s recommended working concentration. Samples dosed with Nedisertib were centrifuged and resuspended in fresh, pre-warmed complete culture media 18-24 hours after dosing. All conditions were run in triplicate.

#### AAV dosing

For KI samples, AAV6 encoding a 1928z anti-CD19 CAR with an EGFRt reporter (Charles River) was added to cells, immediately after RNP delivery at an MOI of 20,000GC cell^-1^, and incubated for 24 hours in serum free culture media at 37°C. After 24 hours, cells were centrifuged to remove AAV and seeded at 10^6^ cells mL^-1^ in complete culture media. For CAR-Ts prepared for the *in vivo* mouse model, cells were centrifuged and resuspended in OptiMEM at 5 × 10^7^ cells mL^-1^. AAV was added at 20,000 MOI and cells were incubated for 1 hour at 37°C. Following the incubation, RNP was added to the cells at the concentration noted above, and processed via Hydroporation. For nucleofection samples, following the 1 hour AAV incubation cells were centrifuged. resuspended in P3 buffer, then RNP was added and the cells were subjected to nucleofection according to the above methods. Following RNP delivery, cells were collected in complete culture media containing 5% human serum.

#### Flow cytometry

Transfected cells at different times post-transfection were analyzed by flow cytometry to measure the cell concentration, viability, KI and KO efficiency. All reagents were used according to manufacturer’s recommendations. To measure viability and cell concentration, cells in media were diluted 1:20 in PBS containing propidium iodide (Sigma-Aldrich, CATt# P4170) then run on the flow cytometer. To measure KI and KO efficiency, cells were pelleted, washed with PBS, and gently resuspended and incubated for 30 min at 4°C in the antibody staining cocktail. The staining cocktail was composed of anti-CD3 FITC (Invitrogen, clone UCHT1) and anti-EGFR APC (Bioloegend, clone AY13) monoclonal antibodies in FACS buffer. After incubation, cells were washed in FACS buffer, pelleted, and resuspended in FACS buffer containing SYTOX Blue viability stain (Thermo Fisher Scientific, CATt# S34857). Samples were then acquired using an Attune NxT flow cytometer (Thermo Fisher Scientific). Compensation was performed using single-stained controls prepared with AbC Total Antibody Compensation Bead Kit (Thermo Fisher Scientific, CAT#A10513). Flow cytometry standard files were exported and analyzed using FlowJo software v3.0 (FlowJo). A standard gating strategy was used to remove debris and aggregated cells. Dead cells were excluded based on SYTOX Blue viability staining.

For identifying T-cell subtypes, flow cytometry was performed on an LSRFortessa X-50 flow cytometer (BD Biosciences). Live lymphocytes were gated based on whether they were wild-type (WT, non-transfected), CAR-(*TRAC* KO only) or CAR+ (*TRAC* KO as well as expressing CAR), as determined by their surface expression of TCRα/β or CAR (G4S scFv linker, Cell Signaling Technology, CAT#69782). From these, we gated on CD4^+^ (BUV395, BD Biosciences,CAT#563550) and CD8^+^ (BV711, BD Biosciences, CAT#569389) T-cells and further divided those into four populations based on their expression of CD45RA (BB515, BD Biosciences, CAT#564552) and CD62L (BV421, BD Biosciences, CAT#563862). Naïve T-cells were defined as CD45RA^+^CD62L^+^, central memory (CM) cells as CD45RA^-^CD62L^+^, effector memory (EM) cells as CD45RA^-^CD62L^-^, and CD45RA^+^ effector memory (EMRA) cells as CD45RA^+^CD62L^-^ cells.

A list of live-dead stain and surface market-targeting antibodies can be found in Supplemental Table 1.

#### Target cells

Firefly luciferase^+^ CD19^+^ NALM6 cells (Imanis CAT#CL150) were cultured in RPMI (Gibco, CAT#11875093) supplemented with FBS (Cytiva CAT#SH30088.03, 10%), sodium pyruvate (Gibco CAT#11360070, 1%), HEPES buffer (Sigma CAT#H0887, 1%), penicillin–streptomycin (Cytiva CAT#SV30010, 1%), non-essential amino acids (Gibco CAT#11140050, 1%) and 2-mercaptoethanol (Gibco CAT#21985023, 0.1%).

#### TCR magnetic depletion

Two days before performing the cytotoxicity assay or infusing into mice, CAR-T populations were enriched by depleting TCRɑ/β+ cells using the Miltenyi human TCRɑ/β depletion kit (Miltenyi CAT#130-133-896). Prior to selection, cell density and viability were assessed using a Cellaca^®^ PLX system (Nexcelom Bioscience). Cells were then centrifuged, resuspended in MACS buffer (80 μL per 10^7^ cells, PBS, 1mM EDTA, 2% Human serum), and incubated with a biotin-conjugated anti-TCRɑ/β antibody (20 μL per 10^7^ cells) for 10 minutes at 4°C. Cells were then washed, resuspended in MACS buffer (80 μL per 10^7^ cells), and incubated with anti-biotin microbeads (20 μL per 10^7^ cells, Miltenyi) for 15 minutes at 4°C. Labeled cells were loaded onto LS Miltenyi MACS columns and processed according to the manufacturer-provided protocol. Cell density in the flow-through from the column was assessed, and isolated cells were centrifuged and resuspended in complete T-cell medium for culture.

#### Cytotoxicity assay

The cytotoxicity of anti-CD19 CAR-T cells was determined by standard luciferase-based assay. In brief, a stable NALM6-Fluc/eGFP cell line served as target cells. The effector (E) and tumor target (T) cells were co-cultured in triplicates at the indicated E:T ratio (1:1 to 1:64) using white-walled 96-well flat clear-bottom plates with 5×10^4^ target cells in a total volume of 100 μL per well in complete T-cell media. The control for maximum signal was NALM6 cells alone, and the control for minimum signal was NALM6 cells and Tween-20 (0.2%). Co-cultures were incubated for approximately 22 h. Then, 100 μL D-luciferin (GoldBio, 0.75 mg ml^−1^) was added to each well, and luminescent signal was measured using a GloMAX Explorer microplate reader (Promega). Cytotoxicity = 100% × (1 − (sample − minimum)/(maximum − minimum)).

#### TIMING^TM^ assays for dynamic single cell analysis

T-Cells were expanded 7 days after transfection and harvested. ≥5×10^6^ CAR-T cells were pelleted and resuspended in cell freezing medium (human serum AB + 5% DMSO) at 1-2 × 10^7^ cells mL^-1^, and frozen in a temperature-controlled freezing unit at -80°C. Frozen cells were thawed in R10 medium: RPMI-1640 (Corning, CAT#10-040-CV), 10%(v/v) dialyzed, heat-inactivated FBS (Hyclone, CAT#SH30079.03), 2 mM L-Glutamine (Corning, CAT#25-005-CI), 1 mM sodium pyruvate (Corning, CAT#4500-710), 20 mM HEPES pH 7.2 (Corning, CAT#25-060-CI), and Penicillin/Streptomycin (50 U mL^-1^, 50 µg mL^-1^, Gibco, CAT#15070-063). After washing and counting, the cells were incubated at 37°C, 5% CO_2_ in R10 containing 60 IU mL^-1^ recombinant human IL-2 (R&D Systems, CAT#202-IL-059/CF). The following day, cells were washed with 1X PBS, blocked in FACS buffer (PBS + 4% FBS v/v) + 5% (v/v) normal goat serum (Sigma, CAT#G9023) for 10 min at 25 °C. Cells were stained with an APC-conjugated anti-human EGFRt antibody (Biolegend, CAT#352905, 10 µg mL^-1^ final concentration) in the same buffer for 30 min on ice, followed by washing with FACS buffer. EGFR^+ve^ (CAR-T) cells were sorted (ABD Biosciences FACSAria, incubated overnight in R10 medium with 60 IU mL^-1^ IL-2 and 100 µg mL^-1^ Normocin™ antibiotic (Invivogen, CAT#ANT-NR-1). The next day, the TIMING assay was performed as previously described^48,50^. Briefly, CAR-T and target T-cells (NALM6, ATCC, CAT#CRL-3273) were separately labeled with fluorescent membrane dyes, PKH67 (Sigma-Aldrich, CAT#PKH67GL-1KT) and PKH26 (Sigma-Aldrich, CAT#PKH26GL-1KT). Labeled cells were pipetted onto nanowell arrays, which were imaged in phenol red-free IMDM medium (Gibco, CAT#21056-023) containing the same supplements as R10 plus AF647-conjugated AnnexinV (Life Technologies, CAT#A23204, 1.6% v/v). Cells were imaged at 5 min intervals over 6 h in a humidified environment at 37°C, 5% CO_2_, as described^47^. CellChorus AI software was used to identify and track cells to quantify multiple parameters, including cell survival, motility, synapse formation and duration, killing, serial killing, and AICD^64^. Data were analyzed using paired Mann-Whitney and t-tests, Kaplan-Meier, and Fisher’s exact tests (GraphPad Prism v10), as detailed in the figure legends.

#### nELISA cytokine Multiplex Assay

To measure secreted cytokine concentration, after an overnight co-culture of CAR-T cells with NALM6 cells (1:1 E:T), 50-100 µL of cell supernatant was collected in a 96 well plate and briefly centrifuged to remove cells and debris. Cleared supernatants were then frozen in a -80°C freezer prior to overnight shipping to Nomic Bio (Montreal, Canada) for subsequent analysis.

Upon arrival, supernatants from treated cells were thawed for nELISA-based secretome analysis using standard protocols, as described previously^65^. Briefly, the nELISA pre-assembles antibody pairs on spectrally encoded microparticles, resulting in spatial separation between non-cognate antibodies, preventing the rise of reagent-driven cross-reactivity, and enabling multiplexing of hundreds of ELISAs in parallel. Protein concentrations on microparticles were read out by high-throughput flow cytometry (Bio-Rad ZE5 cell analyzer) and decoded using Nomic’s proprietary software. Standard curves for all targets were generated to derive pg mL^-1^ values from cytometry fluorescence units. The nELISA MaxPlex panel was used to quantify 187 analytes in each sample.

#### nELISA Cytokine Multiplex Assay Data Analysis

Data analysis was conducted in R (version 4.2.2) with the following packages: tidyverse (v.), data.table (v.), EnhancedVolcano (v.), heatmaply (v.), and reticulate (v.), with associated dependencies within RStudio (v. 2022.12). The data analysis workflow was as follows. Raw values from the nELISA were read into R after reformatting in Microsoft Excel to remove extraneous rows to produce a .csv file containing relevant data rows and the concentration (in picograms per mL) or raw nELISA signal. These were then statistically analyzed for comparisons of interest using one-way ANOVA. The raw signal values were used to calculate the statistical significance, as these values are more dynamic and more closely represent variation in the signal while fold change was calculated based on the transformed concentration values to incorporate data relevant to the underlying biology. The resulting log2fc and p-values were plotted on a volcano plot using the EnhancedVolcano package. These data were also used to generate heatmaps using the heatmaply package by selecting proteins at each condition meeting the following thresholds: ≥|log2fc| and p-value < 0.01. This more stringent p-value was selected to account for multiple comparisons within the data and to reduce the potential for incorrectly calling changes in these proteins. These were then matched to corresponding GO terms, retrieved from Uniprot and DICEDB. Heatmaps were generated using the heatmaply package. These changes were then summarized into bar plots using ggplot by calculating the numbers of proteins meeting these parameters: “up” encompassed proteins with a p-value < 0.01 and a log2fc > 2, “down” included proteins with a p-value < 0.01 and a log2fc <-2, and “same” includes all remaining proteins that fail to meet these criteria.

#### Animal Study

After 2 days of Dynabead activation, T-cells were edited by nucleofection or hydroporation as described above for the KI of a 1928z anti-CD19 CAR with an EGFRt reporter at the *TRAC* locus (M3814 was used for this experiment). TCR depletion was performed 5 days after editing and flow cytometry was conducted subsequently to estimate the CAR%. NOD.Cg-*Prkdc^scid^ Il2rg^tm1Wjl^*/SzJ (NSG) mice were handled ethically and in accordance with the protocol AN182757-01G approved by the University of California, San Francisco (UCSF) Institutional Animal Care and Use Committee. Before and during the experiment, mice were maintained on Clavamox antibiotic. A total of 5 × 10^5^ NALM6 cells were injected into the tail vein of mice that were between 8 and 12 weeks old. After the first BLI measurement (after NALM6 injection, the day before T-cell injection), mice were assigned to each T-cell condition so as to maintain a similar average mass and tumor burden across conditions. Four days after NALM6 injection, 10^5^ or 4 × 10^5^ CAR+ T-cells or an equivalent total number of nucleofected TCR KO T-cells were injected into the tail vein. Mouse health and survival were monitored over time. BLI was performed one or two times per week using a Xenogen in vivo imaging system. At each imaging session, mice were injected intraperitoneally with luciferin (3 mg luciferin per 0.2 ml DPBS) and anesthetized with isoflurane (Medline Industries, CAT#66794-0017-10). The default imaging exposure was 1 min, and shorter exposures were used for images that had a saturating signal at 1 min. Luminescence was quantified using Living Image software (PerkinElmer). Reported BLI values are an average from imaging each mouse on its front and on its back. Mice were euthanized per the approved protocol in the event that they reached end points such as loss of mobility or other signs of morbidity.

## ABBREVIATIONS

4-1BB: Tumor Necrosis Factor Ligand Superfamily Member 9
AAV: Adeno-Associated Virus
AICD: Activation Induced Cell Death
BLI: Bioluminescence Imaging
CAR-T: Chimeric Antigen Receptor T-cell
Cas9: CRISPR-Associated Protein 9
CCL1: Chemokine Ligand 1
CCL5: Chemokine Ligand 5
ccRCC: Clear Cell Renal Cell Carcinoma
CD3: Cluster of Differentiation 3
CD3ζ: Cluster of Differentiation 3 Zeta Chain
CD4: Cluster of Differentiation 4
CD8: Cluster of Differentiation 8
CD19: Cluster of Differentiation 19
CD28: Cluster of Differentiation 28
CD45RA: Cluster of Differentiation 45 RA
CD62L: Cluster of Differentiation 62 L-selectin
CD70: Cluster of Differentiation 70
cGMP: Current Good Manufacturing Practices
CM: Central Memory Cells
CRISPR: Clustered Regularly Interspaced Short Palindromic Repeats
CRS: Cytokine Release Syndrome
CT: Cell Therapy
CXCR1: Cysteine-Amino Acid-Cysteine Motif Chemokine Receptor 1
CXCR2: Cysteine-Amino Acid-Cysteine Motif Chemokine Receptor 2
DIC: Disseminated Intravascular Coagulation
DNA: Deoxyribonucleic Acid
DSB: Double Stranded Break
EGFRt: Truncated Epidermal Growth Factor Receptor
ELISA: Enzyme Linked Immunosorbent Assay
EM: Effector Memory Cells
EMRA: Terminal Effector Memory Cells
E:T: Effector-to-Target
FDA: Food and Drug Administration
G4S: 4-O-Sulfo-Beta-D-Galactopyranose
HDR: Homology Directed Repair
HDRT: Homology Directed Repair Template
HLH: Hemophagocytic Lymphohistiocytosis
Hy-CAR-Ts: Hydroporated Chimeric Antigen Receptor T-cells
ICANS: Immune Effector Cell-Associated Neurotoxicity Syndrome
IL-2: Interleukin 2
IL-6: Interleukin 6
IL-7: Interleukin 7
IL-8: Interleukin 8
IL-10: Interleukin 10
IL-15: Interleukin 15
IL-17A: Interleukin 17A
IL-22: Interleukin 22
IFNγ: Interferon Gamma
KI: Knock-in
KO: Knockout
Log2fc: Logarithmic 2 Fold Change
MHC: Major Histocompatibility Complex
mRNA: Messenger Ribonucleic Acid
NIR: Near Infrared
NSG: NOD.Cg-Prkdc *^scid^* Il2rg *^tm1Wjl^* /SzJ
Nuc-CAR-Ts: Nucleofected Chimeric Antigen Receptor T-cells
P2A: Porcine Teschovirus-1 2A
PBMC: Peripheral Blood Mononuclear Cell
PERC: Peptide-enabled RNP Delivery for CRISPR
RNP: Ribonucleoprotein
ROS: Reactive Oxygen Species
RUO: Research Use Only
scFv: Single-chain Fragment Variable
sgRNA: Single Guide Ribonucleic Acid
TCR (α/β): T-cell Receptor (Alpha/Beta Chains)
TIMING: Time-lapse Imaging Microscopy in Nanowell Grids
TME: Tumor Microenvironment
TNFα: Tumor Necrosis Factor Alpha
TRAC: T-cell Receptor Alpha Constant
Tregs: Regulatory T-cells
WT: Wild Type
μVS: Microfluidic Vortex Shedding

## Acknowledgements

This project has been funded in part with Federal funds from the National Cancer Institute, National Institutes of Health, Department of Health and Human Services, under Contract No. 75N91020C00030 and 7591022C00053. This work was also funded in part by IndieBio (indiebo.co), SOSV (sosv.com), Y Combinator (ycombinator.com), Savantus (savantusventures.com), AusIndustry (business.gov.au), Social Capital (socialcapital.com) Founders Fund’s FF Science (foundersfund.com), Axial (axialsprawl.com), Endpoint Ventures (endpoint.vc), Pioneer Fund (pioneerfund.vc) and BroadOak BioTools Venture Fund (broadoak.com & decibio.com) among others. The UCSF Parnassus Flow Core RRID:SCR_018206 is supported by the DRC Center Grant NIH P30 DK063720. This project was supported by the Cytometry and Cell Sorting Core at Baylor College of Medicine with funding from the CPRIT Core Facility Support Award (CPRIT-RP180672), the NIH (CA125123 and RR024574). This work utilized the nELISA high-throughput protein profiling platform from Nomic Bio (Montreal, Canada). Hydopore^TM^ chips and instruments were manufactured by Microtec (Duisburg, Germany) and Design+Industry (Glebe Island, NSW, Australia), respectively. The authors are thankful for the simulation and computational support provided by Rescale, Amazon Web Services (AWS) and the OpenFOAM community. Certain authors would like to thank Ben Wright, Bryan Poltilove, Geoff Facer and Rich Stoner for their guidance and advice when pursuing science in the startup environment. The content is solely the responsibility of the relevant authors and may not represent the views of anyone acknowledged in this section.

## Author Contributions

B.J.S., V.A., S.B., J.E., R.S.P. conceived and designed the experiments. B.J.S., V.A., S.B., M.F., R.B., F.L.P., A.G.R., W.A.N., Z.L., L.H.W performed the experiments. B.J.S., V.A., S.B., M.F., R.B., L.M.R.F, J.B., L.L., F.L.P., Z.L., L.H.W., J.E., R.S.P. analyzed data. B.J.S., V.A., S.B., F.F., L.H.W., J.E., R.S.P. contributed materials and/or analysis tools. B.J.S., V.A., S.B., F.F., R.B., L.M.R.F, J.E., R.S.P. wrote the paper. All authors read the manuscript and agree with its contents.

## References

1. Mardiana, S., Solomon, B. J., Darcy, P. K. & Beavis, P. A. Supercharging adoptive T cell therapy to overcome solid tumor-induced immunosuppression. Sci Transl Med 11, eaaw2293 (2019).

2. Mardiana, S., Lai, J., House, I. G., Beavis, P. A. & Darcy, P. K. Switching on the green light for chimeric antigen receptor T-cell therapy. Clin Transl Immunology 8, e1046 (2019).

3. Maude, S. L. et al. Chimeric antigen receptor T cells for sustained remissions in leukemia. N Engl J Med 371, 1507–1517 (2014).

4. Porter, D. L., Levine, B. L., Kalos, M., Bagg, A. & June, C. H. Chimeric antigen receptor-modified T cells in chronic lymphoid leukemia. N Engl J Med 365, 725–733 (2011).

5. O’Leary, M. C. et al. FDA Approval Summary: Tisagenlecleucel for Treatment of Patients with Relapsed or Refractory B-cell Precursor Acute Lymphoblastic Leukemia. Clin Cancer Res 25, 1142–1146 (2019).

6. Bouchkouj, N. et al. FDA Approval Summary: Axicabtagene Ciloleucel for Relapsed or Refractory Large B-cell Lymphoma. Clin Cancer Res 25, 1702–1708 (2019).

7. Turtle, C. J. et al. CD19 CAR-T cells of defined CD4+:CD8+ composition in adult B cell ALL patients. J Clin Invest 126, 2123–2138 (2016).

8. Hay, K. A. et al. Kinetics and biomarkers of severe cytokine release syndrome after CD19 chimeric antigen receptor-modified T-cell therapy. Blood 130, 2295–2306 (2017).

9. Wang, X. & Rivière, I. Clinical manufacturing of CAR T cells: foundation of a promising therapy. Mol Ther Oncolytics 3, 16015 (2016).

10. Irving, M., Lanitis, E., Migliorini, D., Ivics, Z. & Guedan, S. Choosing the Right Tool for Genetic Engineering: Clinical Lessons from Chimeric Antigen Receptor-T Cells. Human Gene Therapy 32, 1044–1058 (2021).

11. Zhao, A. et al. Secondary myeloid neoplasms after CD19 CAR T therapy in patients with refractory/relapsed B-cell lymphoma: Case series and review of literature. Front. Immunol. 12. 13, (2023).

12. Brudno, J. N. & Kochenderfer, J. N. Recent advances in CAR T-cell toxicity: Mechanisms, manifestations and management. Blood Rev 34, 45–55 (2019).

13. Chou, C. K. & Turtle, C. J. Assessment and Management of Cytokine Release Syndrome and Neurotoxicity Following CD19 CAR-T Cell Therapy. Expert Opin Biol Ther 20, 653–664 (2020).

14. Morris, E. C., Neelapu, S. S., Giavridis, T. & Sadelain, M. Cytokine release syndrome and associated neurotoxicity in cancer immunotherapy. Nat Rev Immunol 22, 85–96 (2022).

15. Seki, A. & Rutz, S. Optimized RNP transfection for highly efficient CRISPR/Cas9-mediated gene knockout in primary T cells. J Exp Med 215, 985–997 (2018).

16. Eyquem, J. et al. Targeting a CAR to the TRAC locus with CRISPR/Cas9 enhances tumour rejection. Nature 543, 113–117 (2017).

17. Hacein-Bey-Abina, S. et al. Insertional oncogenesis in 4 patients after retrovirus-mediated gene therapy of SCID-X1. J Clin Invest 118, 3132–3142 (2008).

18. Singh, N., Shi, J., June, C. H. & Ruella, M. Genome-Editing Technologies in Adoptive T Cell Immunotherapy for Cancer. Curr Hematol Malig Rep 12, 522–529 (2017).

19. Oh, S. A. et al. High-efficiency nonviral CRISPR/Cas9-mediated gene editing of human T cells using plasmid donor DNA. J Exp Med 219, e20211530 (2022).

20. Schumann, K. et al. Generation of knock-in primary human T cells using Cas9 ribonucleoproteins. Proc Natl Acad Sci U S A 112, 10437–10442 (2015).

21. Muller, Y. D. et al. Precision Engineering of an Anti-HLA-A2 Chimeric Antigen Receptor in Regulatory T Cells for Transplant Immune Tolerance. Front Immunol 12, 686439 (2021).

22. Mandal, P. K. et al. Efficient ablation of genes in human hematopoietic stem and effector cells using CRISPR/Cas9. Cell Stem Cell 15, 643–652 (2014).

23. Gehl, J. Electroporation: theory and methods, perspectives for drug delivery, gene therapy and research. Acta Physiol Scand 177, 437–447 (2003).

24. Chang, A.-Y. et al. Microfluidic Electroporation Coupling Pulses of Nanoseconds and Milliseconds to Facilitate Rapid Uptake and Enhanced Expression of DNA in Cell Therapy. Sci Rep 10, 6061 (2020).

25. Pal, S. K. et al. CD70-Targeted Allogeneic CAR T-Cell Therapy for Advanced Clear Cell Renal Cell Carcinoma. Cancer Discovery OF1–OF14 (2024) doi:10.1158/2159-8290.CD-24-0102.

26. Stadtmauer, E. A. et al. CRISPR-engineered T cells in patients with refractory cancer. Science 367, eaba7365 (2020).

27. Zhang, J. et al. Non-viral, specifically targeted CAR-T cells achieve high safety and efficacy in B-NHL. Nature 609, 369–374 (2022).

28. Thiel, C. & Nix, M. Efficient transfection of primary cells relevant for cardiovascular research by nucleofection. Methods Mol Med 129, 255–266 (2006).

29. Shy, B. R. et al. High-yield genome engineering in primary cells using a hybrid ssDNA repair template and small-molecule cocktails. Nat Biotechnol 41, 521–531 (2023).

30. Peng, W., Polajžer, T., Yao, C. & Miklavčič, D. Dynamics of Cell Death Due to Electroporation Using Different Pulse Parameters as Revealed by Different Viability Assays. Ann Biomed Eng 52, 22–35 (2024).

31. Weaver, J. C. Electroporation of biological membranes from multicellular to nano scales. IEEE Transactions on Dielectrics and Electrical Insulation 10, 754–768 (2003).

32. Hui, S. W. & Li, L. H. In vitro and ex vivo gene delivery to cells by electroporation. Methods Mol Med 37, 157–171 (2000).

33. Meaking, W. S., Edgerton, J., Wharton, C. W. & Meldrum, R. A. Electroporation-induced damage in mammalian cell DNA. Biochimica et Biophysica Acta (BBA) - Gene Structure and Expression 1264, 357–362 (1995).

34. Chicaybam, L., Sodre, A. L., Curzio, B. A. & Bonamino, M. H. An Efficient Low Cost Method for Gene Transfer to T Lymphocytes. PLOS ONE 8, e60298 (2013).

35. Zhang, M. et al. The impact of Nucleofection® on the activation state of primary human CD4 T cells. J Immunol Methods 408, 123–131 (2014).

36. Batista Napotnik, T., Polajžer, T. & Miklavčič, D. Cell death due to electroporation – A review. Bioelectrochemistry 141, 107871 (2021).

37. Dixit, H. G. et al. Massively-Parallelized, Deterministic Mechanoporation for Intracellular Delivery. Nano Lett. 20, 860–867 (2020).

38. Kavanagh, H. et al. A novel non-viral delivery method that enables efficient engineering of primary human T cells for *ex vivo* cell therapy applications. Cytotherapy 23, 852–860 (2021).

39. Loo, J. et al. Microfluidic transfection of mRNA into human primary lymphocytes and hematopoietic stem and progenitor cells using ultra-fast physical deformations. Sci Rep 11, 21407 (2021).

40. Sido, J. M. et al. Electro-mechanical transfection for non-viral primary immune cell engineering. 2021.10.26.465897 Preprint at 10.1101/2021.10.26.465897 (2021).

41. Graham, C., Jozwik, A., Pepper, A. & Benjamin, R. Allogeneic CAR-T Cells: More than Ease of Access? Cells 7, 155 (2018).

42. Abou-el-Enein, M. et al. Scalable Manufacturing of CAR T Cells for Cancer Immunotherapy. Blood Cancer Discov 2, 408–422 (2021).

43. Jarrell, J. A. et al. Intracellular delivery of mRNA to human primary T cells with microfluidic vortex shedding. Sci Rep 9, 3214 (2019).

44. Jarrell, J. A. et al. Numerical optimization of microfluidic vortex shedding for genome editing T cells with Cas9. Sci Rep 11, 11818 (2021).

45. Foss, D. V. et al. Peptide-mediated delivery of CRISPR enzymes for the efficient editing of primary human lymphocytes. *Nat*. Biomed. Eng 7, 647–660 (2023).

46. Merouane, A. et al. Automated profiling of individual cell–cell interactions from high-throughput time-lapse imaging microscopy in nanowell grids (TIMING). Bioinformatics 31, 3189–3197 (2015).

47. Lu, H. et al. TIMING 2.0: high-throughput single-cell profiling of dynamic cell-cell interactions by time-lapse imaging microscopy in nanowell grids. Bioinformatics 35, 706–708 (2019).

48. Romain, G. et al. Multidimensional single-cell analysis identifies a role for CD2-CD58 interactions in clinical antitumor T cell responses. J Clin Invest 132, (2022).

49. Rezvan, A. et al. Identification of a clinically efficacious CAR T cell subset in diffuse large B cell lymphoma by dynamic multidimensional single-cell profiling. Nat Cancer (2024) doi:10.1038/s43018-024-00768-3.

50. Liadi, I. et al. Defining potency of CAR+ T cells: Fast and furious or slow and steady. Oncoimmunology 8, e1051298 (2019).

51. Melenhorst, J. J. et al. Decade-long leukaemia remissions with persistence of CD4+ CAR T cells. Nature 602, 503–509 (2022).

52. Blaeschke, F. et al. Modular pooled discovery of synthetic knockin sequences to program durable cell therapies. Cell 186, 4216–4234.e33 (2023).

53. Pavlovic, K. et al. Generating universal anti-CD19 CAR T cells with a defined memory phenotype by CRISPR/Cas9 editing and safety evaluation of the transcriptome. Front. Immunol. 15, (2024).

54. Morshedi Rad, D., et al. A Comprehensive Review on Intracellular Delivery. Advanced Materials 33, 2005363 (2021).

55. Stewart, M. P., Langer, R. & Jensen, K. F. Intracellular Delivery by Membrane Disruption: Mechanisms, Strategies, and Concepts. Chem. Rev. 118, 7409–7531 (2018).

56. Bickel, M. The role of interleukin-8 in inflammation and mechanisms of regulation. J Periodontol 64, 456–460 (1993).

57. Jin, L. et al. CXCR1- or CXCR2-modified CAR T cells co-opt IL-8 for maximal antitumor efficacy in solid tumors. Nat Commun 10, 4016 (2019).

58. Zenewicz, L. A. IL-22: There Is a Gap in Our Knowledge. ImmunoHorizons 2, 198–207 (2018).

59. Tosti, N. et al. Infiltration by IL22-Producing T Cells Promotes Neutrophil Recruitment and Predicts Favorable Clinical Outcome in Human Colorectal Cancer. Cancer Immunology Research 8, 1452–1462 (2020).

60. Mei, Z. et al. MUC1 as a target for CAR-T therapy in head and neck squamous cell carinoma. Cancer Med 9, 640–652 (2020).

61. Trinchieri, G. Interleukin-10 production by effector T cells: Th1 cells show self control. J Exp Med 204, 239–243 (2007).

62. Jiang, Y., Li, Y. & Zhu, B. T-cell exhaustion in the tumor microenvironment. Cell Death Dis 6, e1792–e1792 (2015).

63. Seif, F., Torki, Z., Zalpoor, H., Habibi, M. & Pornour, M. Breast cancer tumor microenvironment affects Treg/IL-17-producing Treg/Th17 cell axis: Molecular and therapeutic perspectives. Molecular Therapy - Oncolytics 28, 132–157 (2023).

64. Teachey, D. T. et al. Identification of Predictive Biomarkers for Cytokine Release Syndrome after Chimeric Antigen Receptor T-cell Therapy for Acute Lymphoblastic Leukemia. Cancer Discovery 6, 664–679 (2016).

65. Dagher, M. et al. nELISA: A high-throughput, high-plex platform enables quantitative profiling of the secretome. 2023.04.17.535914 Preprint at 10.1101/2023.04.17.535914 (2023).

