## Supplemental Tables and Figures for "Scalable intracellular delivery via microfluidic vortex shedding enhances the function of chimeric antigen receptor T-cells"

### Contents

#### Supplemental figures


1. Computational fluid dynamics simulations and related biological data for different flow cell widths - Supplemental Fig. 1
2. Representative flow cytometry gating for assessing TCR surface expression following Cas9 TRAC-RNP delivery to and CAR AAV transduction of CD3+ T cells, TCR KO% and KIKO ratios - Supplemental Fig. 2
3. Representative flow cytometry gating for assessing CD62L/CD45RA phenotypes.- Supplemental Fig. 3
4. Additional single-cell TIMING data for the five donors - Supplemental Fig. 4
5. Bioluminescence imaging trace for each mouse - Supplemental Fig. 5
6. Representative flow cytometry gating for TCR/EGFR surface expression before and after magnetic TCR depletion ahead of *in vitro* NALM6 co-culture or infusion into mice. - Supplemental Fig. 6


#### Supplemental tables


1. Antibodies used for flow cytometry analysis - Supplemental Table 1


#### Supplemental videos


1. Montage and video of individual time points (hh:mm) during a TIMING assay - Supplemental Video 1

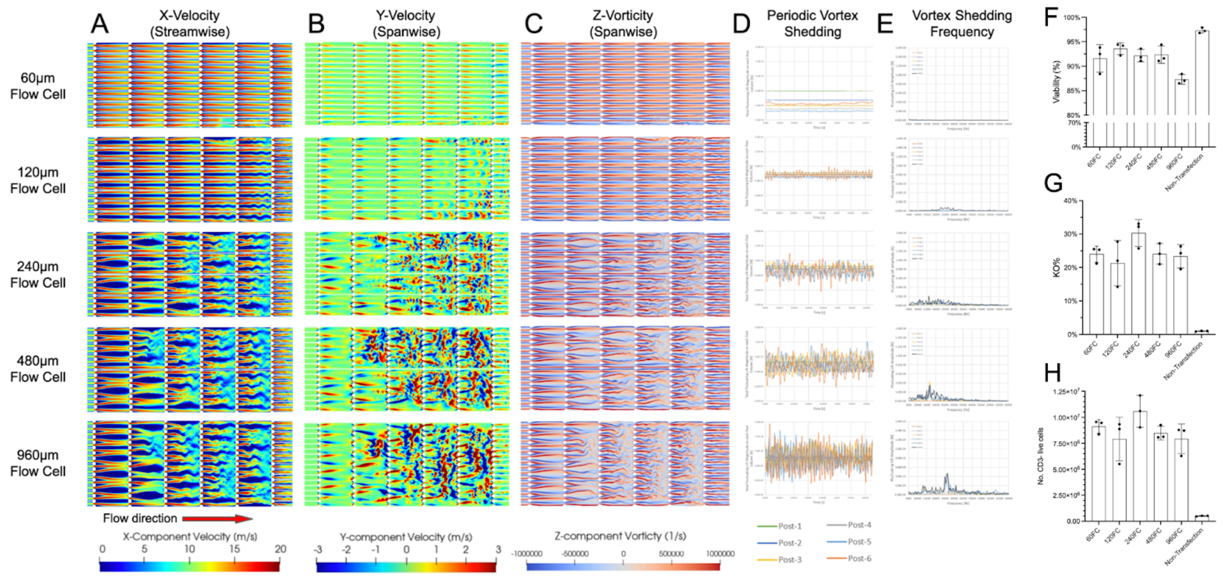


**Supplemental Fig. 1 |** Computational fluid dynamics simulations and related biological data for different flow cell widths showing **A)** x-velocity, **B)** y-velocity, **C)** z-vorticity, **D)** time-domain force acting on individual posts, **E)** frequency-domain force acting on individual posts, **F)** cell viability at 24 hr, **G)** TRAC-1 KO efficiency on day 7, and **H)** TRAC-1 KO yield on day 7. All data points for f-h involve n = 3 biological data points and the error bars represent 1 standard deviation. Optimal 240 µm flow cell width equivalent to 1x flow cell and data related to Fig 2a,b.

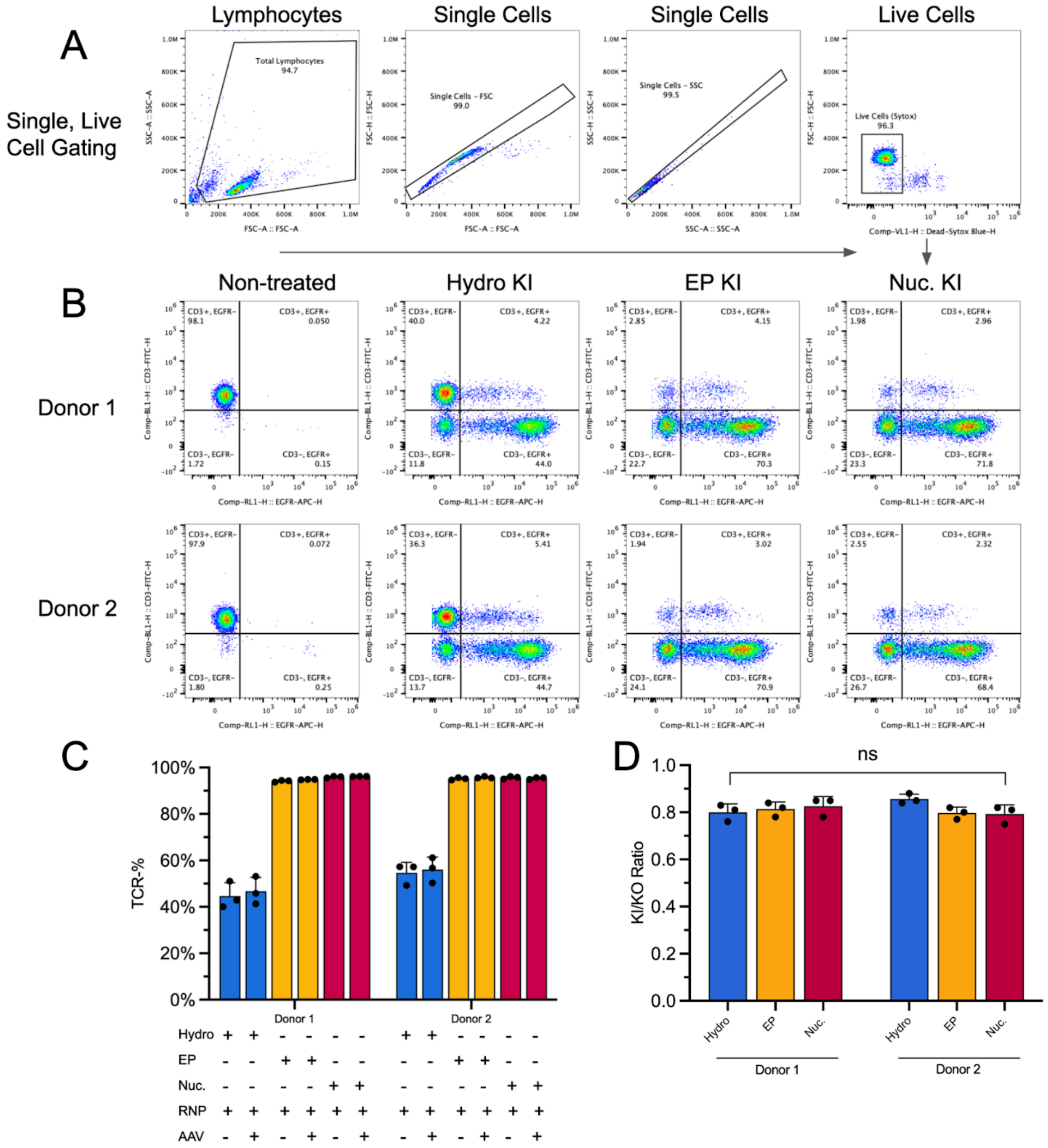


**Supplemental Fig. 2 |** Representative flow cytometry gating for assessing TCR surface expression following Cas9 TRAC-RNP delivery to and CAR AAV transduction of CD3+ T cells for **A)** Singe, live cell gating strategy **B)** Representative TCR KO/EGFR KI flow plots for three different transfection methods. Data are related to Fig 1g,h **C)** TCR knock-out efficiencies for the three different transfection methods +/- the addition of the AAV HDRT. **D)** The ratio of knock-in efficiency to knockout efficiency. Statistical significance by Brown-Forsythe ANOVA test, n=3, p= 0.32.

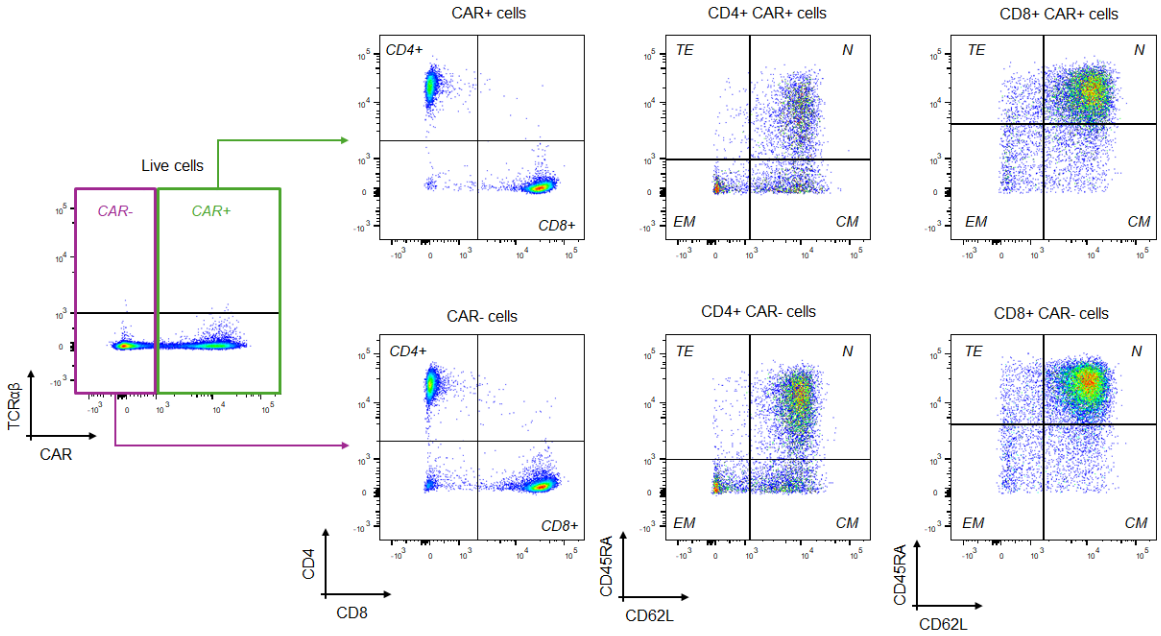


**Supplemental Fig. 3 |** Representative flow cytometry gating for assessing CD62L/CD45RA phenotypes in CD4+ and CD8+ T cells. Phenotypes were assessed independent of editing outcome. Data related to Fig 3a.

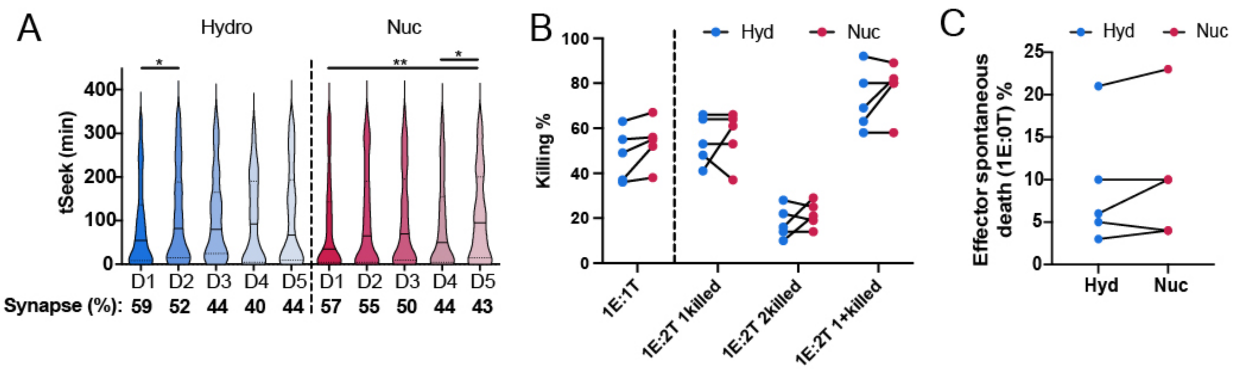


**Supplemental Fig. 4 |** Additional single-cell TIMING data for the five donors shown in Fig. 4. **A)** tSeek (time for CAR-T cells to form a synapse with a target cell). The percentage of CAR-Ts that formed a synapse with one target (1E:1T nanowells) is shown below the x-axis. **B)** Percent of CAR-Ts that formed a synapse and killed one or more targets after synapse formation. **C)** 0T nanowells (1 CAR-T: 0 target cells = no target cells present) that were dead after the six-hour assay. Statistical tests were performed and denoted as described in Fig. 4.

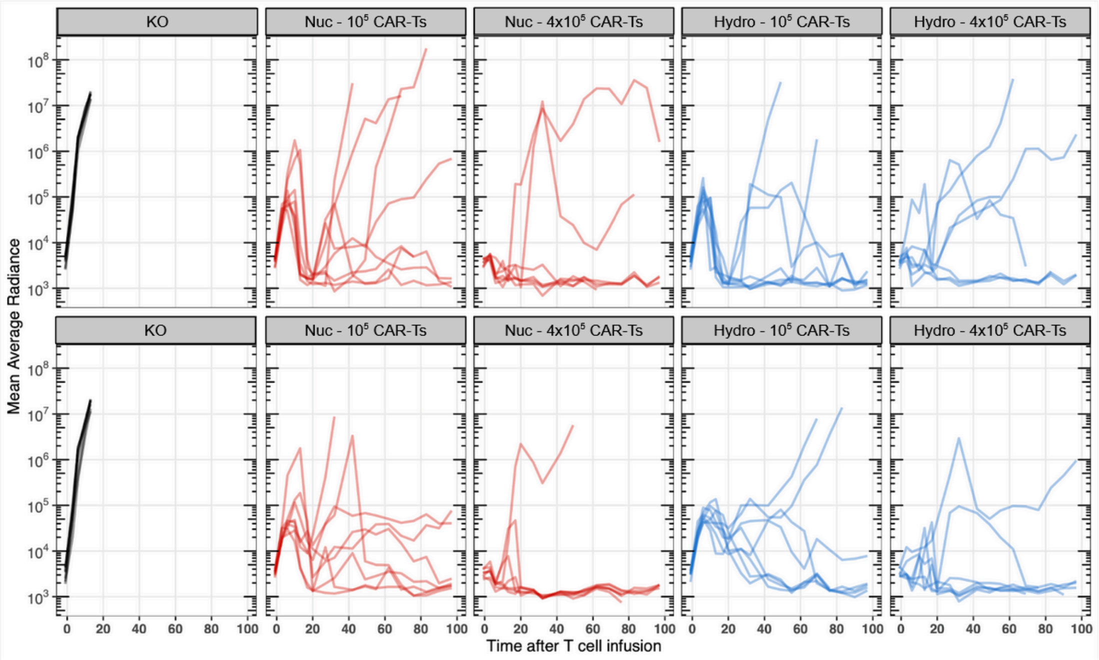


**Supplemental Fig. 5 |** Bioluminescence imaging trace for each mouse associated with data in Fig. 5c.

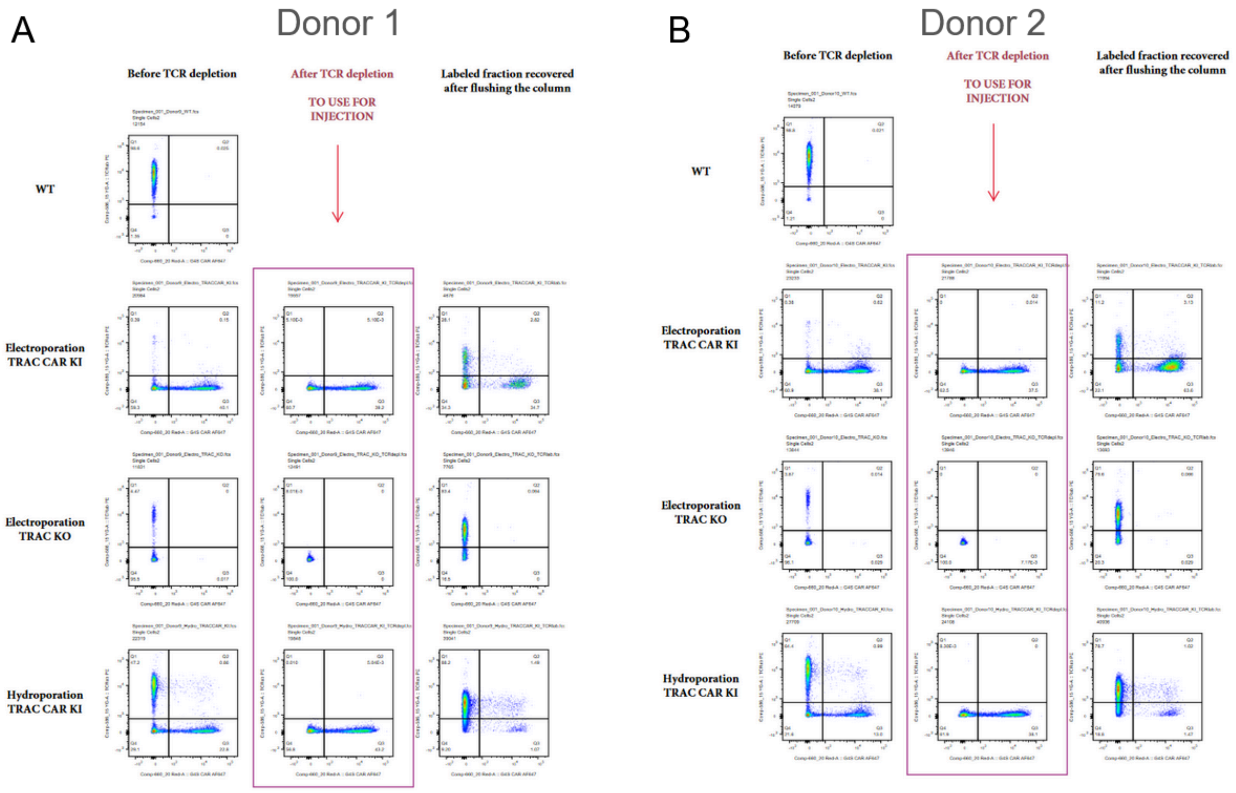


**Supplemental Fig. 6 | ** Representative flow cytometry gating for assessing TCR/EGFR surface expression following Cas9 TRAC-RNP delivery to and CAR AAV transduction of CD3+ T cells for **A)** Donor 1 and **B)** Donor 2. Data are related to Fig 5.

**Supplemental Table. 1 |** Antibodies/stains used for flow cytometry analysis.

| Target | Fluorophore | Dilution | Manufacturer | Clone | Catalog # |
|---:|---:|---:|---:|---:|---:|
| CD3 | AF700 | 1:100 | Thermo Fisher Scientific | UCHT1 | 56-0038-42 |
| CD3 | FITC | 1:100 | BioLegend | UCHT1 | 300405 |
| CD4 | PE-Cy7 | 1:500 | BioLegend | OKTA | 317413 |
| CD8 | PE-Texas Red | 1:500 | Thermo Fisher Scientific | 3B5 | MHCD0817 |
| PD-1 | APC | 1:200 | Thermo Fisher Scientific | J105 | 17-2799-41 |
| Lag3 | PE | 1:100 | Thermo Fisher Scientific | 3DS223H | 12-2239-41 |
| CD27 | FITC | 1:100 | Thermo Fisher Scientific | O323 | 11-0279-42 |
| CD197 (CCR7) | APC-eF780 | 1:200 | Thermo Fisher Scientific | 3D12 | 47-1979-42 |
| CD45RA | PerCP-Cy5.5 | 1:200 | Thermo Fisher Scientific | H100 | 45-0458-41 |
| TCRα/β | FITC | 1:200 | Miltenyi Biotec | REA652 | 130-113-538 |
| EGRP | APC | 1:200 | BioLegend | AY13 | 352906 |
| n/a | GhostDye Red 780 | 1:1000 | Tonbo Biosciences | n/a | 13-0865-T500 |
| n/a | Sytox Blue | 1:1000 | Thermo Fisher Scientific | n/a | S34857 |
| n/a | Propidium Iodide (PI) | 1:1000 | Thermo Fisher Scientific | n/a | P4170 |

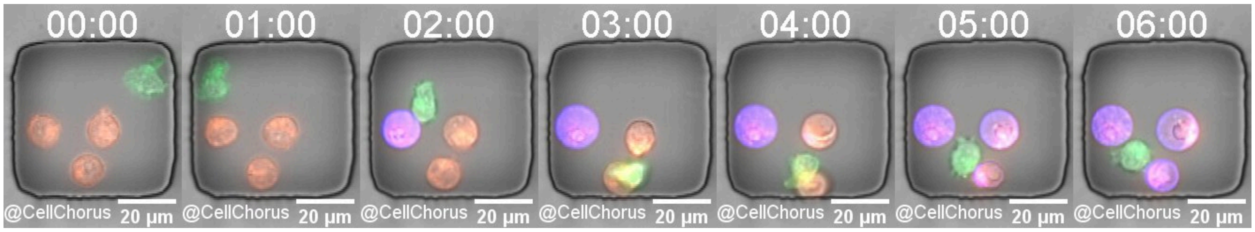


**Supplemental Video. 1 |** Montage and video of individual time points (hh:mm) during a TIMING assay of Hyd-CAR-T (green) interacting with three NALM6 target cells (red) in a nanowell. Apoptotic cells are identified with AnnexinV-Alexa674 (purple) during the assay. This Hyd-CAR-T is a serial killer and killed all three target cells in the nanowell within 6 hrs.
